# NDR kinase SAX-1 controls dendrite branch-specific elimination during neuronal remodeling in *C. elegans*

**DOI:** 10.1101/2025.06.09.658633

**Authors:** Paola V. Figueroa-Delgado, Shaul Yogev

**Affiliations:** Department of Cell Biology, Yale School of Medicine, 295 Congress Ave, New Haven, CT 05610; Department of Neuroscience, Yale School of Medicine, 100 College St, New Haven, CT 06510

**Keywords:** neuronal remodeling, dendrite pruning, dauer, SAX-1/NDR, SAX-2/Fry, MOB-2/Mob, membrane trafficking, RABI-1/Rabin8, small GTPase RAB-11.2

## Abstract

Neuronal remodeling is crucial for proper nervous system development and function, and can be initiated by developmental programs, activity-dependent mechanisms, or stress. Despite significant advances, the underlying mechanisms that govern this process remain poorly understood. Here, we adapted *C. elegans* IL2 sensory dendrites as a model system to study developmental and stress-mediated dendrite pruning. Upon entering a stress-induced developmental diapause, IL2 dendrites grow a complex dendritic arbor, which is later pruned when reproductive development resumes. We identified unexpected specificity in the pruning process, with distinct genetic requirements to direct branch-specific elimination of secondary, tertiary, and quaternary branches. The serine/threonine kinase SAX-1/NDR promotes elimination of secondary and tertiary, but not quaternary, dendrites. SAX-1 functions with its conserved interactors SAX-2/Furry and MOB-2 in the removal of both dendritic branches. The guanine-nucleotide exchange factor RABI-1/Rabin8 and the small GTPase RAB-11.2 mediate the elimination of secondary branches with SAX-1, but their effect on tertiary branches is minimal. Consistent with the known roles of RABI-1 and RAB-11.2 in regulating membrane dynamics, we find that SAX-1 promotes endocytosis during remodeling. Together, our findings reveal distinct mechanisms for branch-specific elimination under stress-induced and developmentally regulated neuronal remodeling.

## Introduction

Neuronal remodeling is essential for proper nervous system development and function. Dendrite remodeling can be initiated by stereotyped developmental programs, activity-dependent mechanisms or stress (Brunson et al. 2005; Schuldiner and Yaron 2015; Riccomagno and Kolodkin 2015; McEwen, Nasca, and Gray 2016; Furusawa and Emoto 2021). For example, in *Drosophila,* dendrite pruning is regulated by developmental ecdysone signaling in the mushroom body and in peripheral sensory neurons during morphogenesis (Lee et al. 2000; Kuo, Jan, and Jan 2005). In contrast, in mammalian hippocampal neurons, chronic stress has been shown to promote dendrite retraction (Magariños et al. 1996; Vyas et al. 2002; Christian et al. 2011; McEwen, Nasca, and Gray 2016). Although deregulated neuronal remodeling has been suggested to be involved in neurodevelopmental and neuropsychiatric disorders such as Autism Spectrum Disorder and Schizophrenia (Riccomagno and Kolodkin 2015), the mechanisms that govern this process remain poorly understood.

A myriad of cell biological mechanisms cooperate to direct dendrite pruning, including protease activation, transport, cytoskeletal dynamics, and membrane dynamics (Williams and Truman 2005; Kuo et al. 2006; Lin et al. 2015; Riccomagno and Kolodkin 2015; Schuldiner and Yaron 2015; Krämer, Rode, and Rumpf 2019; Rumpf, Wolterhoff, and Herzmann 2019; Furusawa and Emoto 2021; Rui 2024). Evidence for the role of membrane dynamics comes primarily from *Drosophila* class IV da neurons, where localized endocytic events at proximal dendrites correlate with membrane thinning and precede pruning of the dendritic arbor (Kanamori et al. 2013; H. Zhang et al. 2014; Kanamori et al. 2015). In the same system, the recycling endosome protein Rab11 is required for dendrite pruning, at least partially through the removal of the cell surface protein Neuroglian (H. Zhang et al. 2014; Krämer, Rode, and Rumpf 2019; Lin et al. 2020). Rab11 can bind the <u>g</u>uanine-nucleotide <u>e</u>xchange factor (GEF) Rabin8, which can activate Rab8 and Rab10 (Knödler et al. 2010; Westlake et al. 2011; Feng et al. 2015; Homma and Fukuda 2016). Although Rab11 has been implicated in membrane retrieval and removal of surface Neuroglian, whether Rabin8, and its associated GTPases, function in dendrite pruning has not been investigated.

<u>N</u>uclear <u>D</u>bf2-related (NDR) kinases are AGC family serine/threonine kinases that are evolutionarily conserved from yeast to humans (Tamaskovic, Bichsel, and Hemmings 2003; Hergovich et al. 2006; Santos et al. 2023). NDR kinases regulate cell shape, growth, and polarity (Verde, Wiley, and Nurse 1998; Zallen et al. 2000; Geng et al. 2000; Hergovich et al. 2006; Chen et al. 2019). Mutations in NDR kinases lead to ectopic membrane growth in *C. elegans* and mammalian neurons (Gallegos and Bargmann 2004; Zallen et al. 2000; Roşianu et al. 2023), and to excessive dendrite branching in *Drosophila*, consistent with a role in restricting cellular growth (Emoto et al. 2004). The mechanism by which NDR kinases restrict cellular growth remains unclear, although several NDR substrates, including Rabin8, regulate membrane dynamics (Ultanir et al. 2012; Roşianu et al. 2023). Whether NDR kinases, in addition to restricting neurite growth, promote neurite elimination is unknown.

Here, we establish *C. elegans* inner labia 2 dorsal and ventral (IL2Q) neurons as a genetically tractable model for dendrite pruning. IL2Q primary dendrites elaborate new branches when *C. elegans* enters a stress-induced and developmentally encoded quiescence-state known as dauer arrest (Schroeder et al. 2013; Androwski, Flatt, and Schroeder 2017). These newly generated branches are then eliminated upon return to favorable conditions (Schroeder et al. 2013). We find that IL2Q pruning shows branch-specific and stress-specific genetic requirements: the NDR kinase SAX-1 is required for pruning secondary and tertiary, but not quaternary branches. SAX-1 promotes branch elimination in post dauer larvae induced by the *daf-7/*TGF-β or *daf-2/*Insulin-receptor pathways, but not by starvation. SAX-1 functions with its conserved interactors SAX-2/Furry and MOB-2 to regulate membrane dynamics during IL2Q pruning. SAX-1 functions with the guanine-nucleotide exchange factor RABI-1/Rabin8 and the small GTPase RAB-11.2, which are primarily required for secondary dendrite elimination. These results provide insights into cell-biological mechanisms that underlie remodeling of complex dendritic arbors during development and stress.

## Results

### *shy87* mutants disrupt dendrite remodeling

To study dendrite remodeling we adapted the Inner <u>L</u>abial 2 (IL2) sensory neurons of *C. elegans* as an experimental model. IL2s extend an anterior primary dendrite that terminates in a sensory cilium at the tip of the nose, and project short axons that run posteriorly to the nerve ring, where they turn circumferentially (White et al. 1986). The IL2 dendrite is unbranched in well-fed animals undergoing reproductive development. However, two IL2 pairs (IL2D and IL2V, together referred to as IL2Q) undergo extensive and stereotypic branching during dauer arrest – an alternative developmental diapause that is induced by stress conditions such as overpopulation or starvation (**Figure 1A, B**). In dauer, IL2Q primary (1°) dendrites extend secondary (2°), tertiary (3°), and quaternary (4°) branches at roughly 90° to each other, forming a characteristic dendrite arbor (**Figure 1C; Supplemental Figure 1A**). Upon re-exposure to favorable conditions, IL2Q eliminate most of their dauer-generated dendritic branches (**Figure 1B; 1D; Supplemental Figure 1A-B**), leaving primary dendrites intact (Schroeder et al. 2013). The factors that control IL2Q dendrite elimination following dauer recovery are unknown.

**Figure 1.**
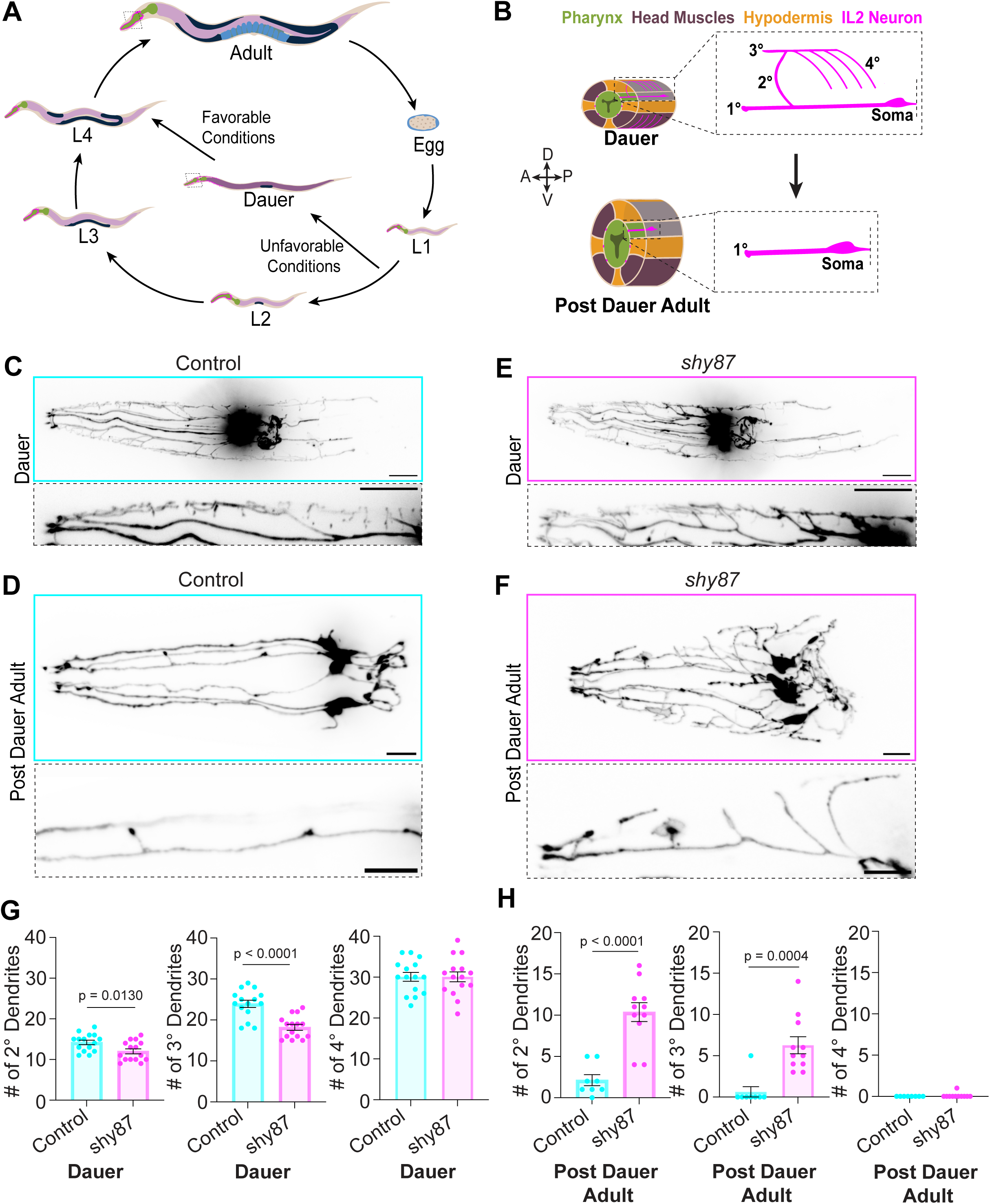
*shy87* mutants exhibit a remodeling defect in post dauer adults. (A) Schematic depicting *C. elegans* life cycle. Adapted from WormAtlas. Under unfavorable conditions, the nematode arrests into an alternative developmental molt (dauer). Upon re-exposure to favorable conditions, reproductive development is resumed into adulthood. (B) Oblique transverse schematic of IL2Q neurons (magenta) at dauer arrest (top) and in a post dauer adult worm (bottom). Pharynx (green); Head muscles (purple); Hypodermis (yellow). At dauer, the IL2Q neurons extend a stereotypical dendritic arbor characterized by a 2° dendrite extending from a 1° dendrite towards the dorsal midline; 3° dendrites bifurcate from 2° dendrite towards the anterior and posterior in parallel to 1° dendrite; 4° dendrites extending into the head muscle quadrants (purple). When reproductive development is resumed, post dauer adult worms remove the higher-order dendrites, leaving the 1° dendrite intact. (C-D) Z-projection of control dauer (C) and post dauer adult (D) expressing *tba-6*p::tagRFP in IL2s (top). Zoomed inset of select IL2 dorsal neuron (bottom, dashed box). (E-F) Maximum intensity Z-projection of *shy87* mutant dauer (E) and post dauer adult (F) expressing cytosolic tagRFP under *tba-6*p (top). Zoomed inset of select IL2 dorsal neuron (bottom, dashed box). Scale bar, 10μm. (G) Quantification of the number of IL2Q 2°-4° dendrites at dauer comparing *shy87* mutants (n = 15) to control animals (n = 15). Unpaired *t*-test. (H) Quantification of the number of IL2Q higher-order dendrites in control (n = 8) versus *shy87* mutants post dauer adults (n = 11). Unpaired *t*-test with Welch’s correction (2°) or Mann-Whitney U test (3°-4°). Error bars are ±SEM.

To uncover new regulators of neuronal remodeling, we first optimized conditions for an unbiased visual genetic screen for IL2Q remodeling-defective mutants. IL2 neurons were visualized by expressing a cytosolic TagRFP driven by the *tba-6* promoter, an α-tubulin isoform that is enriched in IL2s (Hurd et al. 2010; Schroeder et al. 2013; Nishida et al. 2021). We synchronized dauer entry and exit using a temperature-sensitive allele of the TGF-β homolog *daf-7(e1372),* which causes constitutive dauer entry at 25°C and allows recovery from dauer and the resumption of reproductive development at 15°C (Riddle, Swanson, and Albert 1981; Ren et al. 1996). IL2Q morphology at dauer and in post dauer adults was qualitatively similar in *daf-7(e1372)* and wildtype N2 animals (**Supplemental Figure 1D**). Hence, unless otherwise noted, we use *daf-7(e1372)* as a control strain.

From a screen of roughly 3,000 haploid genomes (screen strategy outlined in **Supplemental Figure 1C)**, we isolated a mutant, *shy87*, that exhibited a striking maintenance of IL2Q dendrite branches as post dauer adults. *shy87* mutants eliminated 4° dendritic branches but showed excessive 2° and 3° dendritic branches compared to control (**Figure 1C-H**), suggesting that *shy87* is required for the elimination of secondary and tertiary dendritic branches. To test whether this defect reflects a failure to eliminate dauer-generated branches or an earlier developmental defect, we quantified the total number of IL2Q dendrites across 2°-4° branch orders in control and *shy87* mutants at dauer (**Figure 1G**). *shy87* mutant dauers showed a minor reduction in secondary and tertiary branches compared to control (**Figure 1G**). These results indicate that *shy87* is specifically required for the elimination of dauer-generated dendrite branches.

### The conserved serine/threonine kinase *sax-1/NDR* is required for dendrite branch-specific elimination

We next used whole-genome sequencing and SNP mapping to identify a signature <u>E</u>thyl <u>M</u>ethane<u>s</u>ulfonate (EMS) mutation (G>A) that converts a glycine to an asparagine at position 316 of the conserved serine/threonine kinase SAX-1 in *shy87* mutants (**Figure 2A**). We validated *sax-1(shy87)* as the causal mutation for pruning defects with three complimentary approaches. First, we tested a partial deletion allele, *sax-1(ky491),* and found that it phenocopied *shy87* (**Figure 2E-F**). Second, we generated an early stop codon in *sax-1* in control animals using CRISPR-Cas9, which led to pruning defects indistinguishable from *shy87* (**Figure 2C;2E-F**). Lastly, we reverted the mutated asparagine 316 back to glycine in *shy87* mutants and found that this rescued the mutant phenotype (**Figure 2D-F**). Together, these results indicate that *shy87* is a mutation in *sax-1,* and that *sax-1* is required for IL2Q dendrite remodeling.

**Figure 2.**
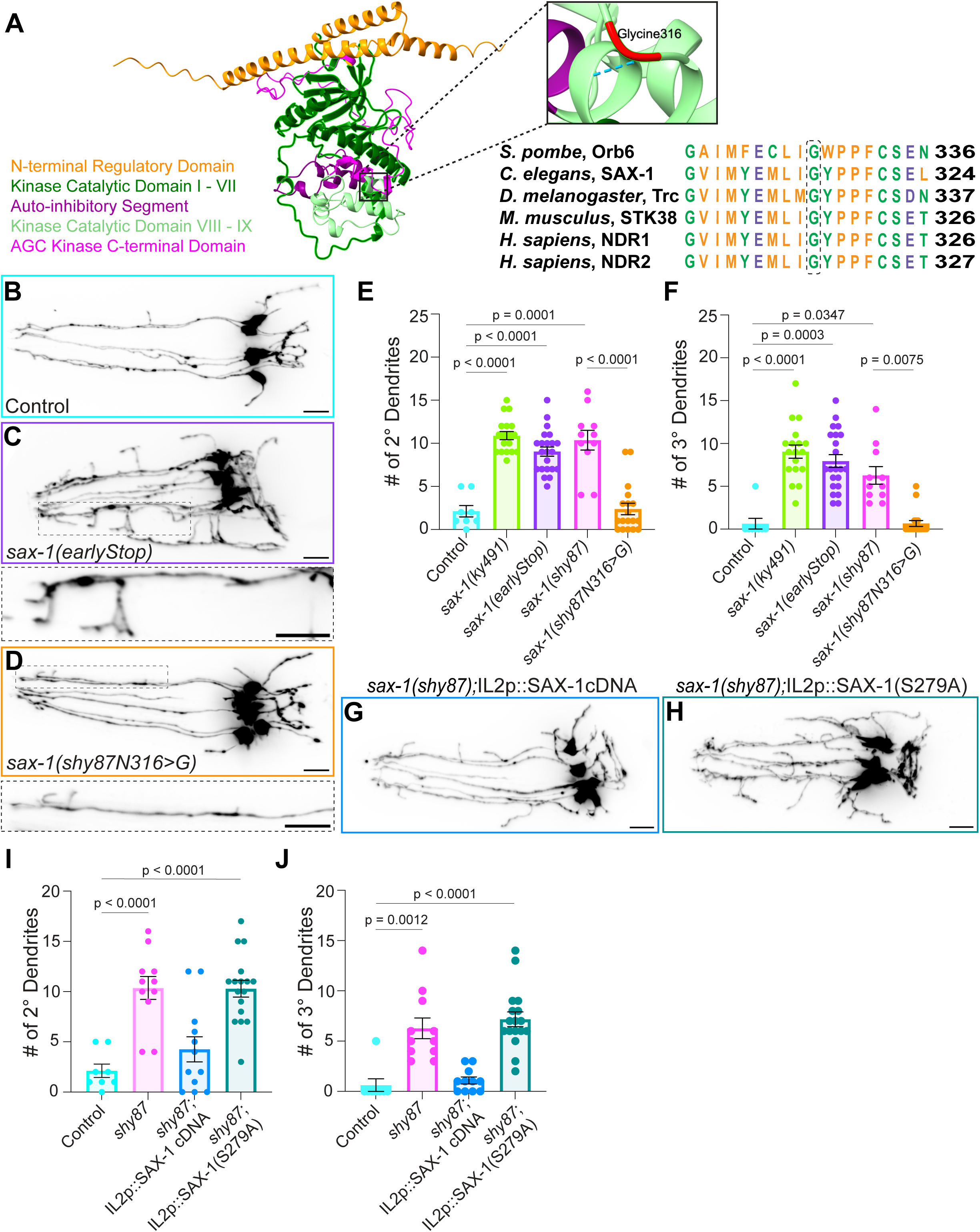
*SAX-1/NDR* promotes IL2Q remodeling in a cell-specific and kinase activity-dependent manner. (A) AlphaFold predicted structure of SAX-1 with key functional domains indicated: N-terminal regulatory domain (1-79AA, orange), kinase catalytic domain I-VII (80-270AA, dark green), auto-inhibitory sequence (270-310AA, dark purple), kinase catalytic domain VIII-IX (310-370AA, light green), and the AGC family kinase C-terminal domain (370-446AA, magenta). Close-up view of glycine316 (red), which is conserved across species from yeast to human homologs of NDR1/2 kinases and mutated into an asparagine in *shy87* mutants. (B-D) Maximum intensity Z-projection of control (B), early stop (C), and engineered repair of the *shy87* allele (D) post dauer adult animals, demonstrating that loss of *sax-1* function is responsible for the pruning defects in *shy87* mutants. Zoomed inset of select IL2Q are shown for *early stop* (C) and *shy87* engineered repair (D) with a dashed box. Scale bars, 10μm. (E, F) Quantifications of IL2Q 2° dendrite (E) and 3° dendrite number in post dauer adults of control and indicated *sax-1* alleles. Brown-Forsythe and Welch ANOVA with Dunnett’s correction (E) or Kruskal-Wallis test with Dunn’s correction (F), all genotypes compared to control and *shy87* mutants, n = 8-22. All *sax-1* mutant alleles phenocopied *shy87.* Repair of the *shy87* missense point mutation rescued the mutant phenotype back to control numbers. (G-H) Maximum intensity Z-projection of *shy87* post dauer adult mutants expressing wildtype SAX-1 cDNA (G) and a kinase-dead (S279) construct (H) under an IL2-specific promoter (*tba-6*p). Scale bars, 10μm. (I, J) Quantifications comparing the total number of IL2Q 2° dendrites (I) and 3°dendrites (J) in post dauer adults of control, *sax-1* mutants, and indicated rescue transgenes. Brown-Forsythe and Welch ANOVA with Dunnett’s multiple comparison (I) and Kruskal-Wallis with Dunn’s correction (J), n = 8-21. Error bars are ±SEM.

NDR kinases are required to restrict cell size across organisms (Tamaskovic, Bichsel, and Hemmings 2003; Gallegos and Bargmann 2004; Emoto et al. 2004; Hergovich et al. 2006; Yd et al. 2019; Roşianu et al. 2023). In *C. elegans* and mice, NDR mutants show excessive growth of neuronal membranes, and in *Drosophila,* mutations in the NDR homolog Tricornered lead to excessive dendrite branching and loss of self-avoidance (Zallen et al. 2000; Gallegos and Bargmann 2004; Emoto et al. 2004; Roşianu et al. 2023). It is therefore unexpected that in IL2Q *sax-1* is required to eliminate existing processes rather than to restrict growth. To confirm that dendrite branch-maintenance in *sax-1* mutants is not caused by ectopic regrowth after initial elimination, we systematically characterized the time course of dendrite remodeling following recovery from dauer arrest. In control animals, 4° dendrites were mostly eliminated by 16 hours. Most 3° branches were eliminated between 16-22 hours (**Supplemental Figure 2A**), whereas 2° branches were eliminated closer to the L4 molt. Examination of *sax-1* mutants at 16-20 hours, when most 3° branches undergo elimination, revealed that these branches are maintained in the mutants (**Supplemental Figure 2B**). This result strongly argues that excessive dendrite branches in *sax-1* mutant adults reflect a failure to eliminate dauer-born branches rather than ectopic branch regrowth.

Dauer entry involves drastic changes to animal physiology (Cassada and Russell 1975; Riddle 1977; Golden and Riddle 1984; Gerisch et al. 2001; Burnell et al. 2005). Furthermore, the method of dauer induction can affect the characteristics of the dauer larvae (Golden and Riddle 1982; Karp 2018; Hu 2018). We therefore asked whether the requirement for *sax-1* in controlling IL2Q dendrite morphology depends on the selected method of dauer induction. We found that *sax-1* mutants that have not undergone dauer arrest (*i.e.* normal development) exhibit normal IL2Q morphology, suggesting that *sax-1* only plays a role post dauer (data not shown). Next, we tested whether the requirement for *sax-1* is specific to dauers induced by the temperature-sensitive *daf-7*/TGF-β mutation. For this, we examined IL2Q morphology in *sax-1* mutants following dauer arrest induced by starvation or by using a mutant allele of the insulin receptor *daf-2(e1370)* (**Supplemental Figure 2C**), which promotes dauer entry through a different pathway than *daf-7(e1372)* (Riddle, Swanson, and Albert 1981; Gottlieb and Ruvkun 1994; Ren et al. 1996; Hu 2018; Karp 2018). Interestingly, starved animals did not require *sax-1* for dendrite remodeling, whereas in *daf-2* mutants, similar to *daf-7* mutants, *sax-1* was required for dendrite remodeling (**Supplemental Figure 2C-D**). These results suggest that the requirement for *sax-1* in IL2Q dendrite morphology is not universal but relies on specific physiological states of the dauer stage.

### SAX-1/NDR acts cell-autonomously and is kinase activity dependent

To determine whether SAX-1 functions cell autonomously in IL2Q to promote elimination, we expressed *sax-1* cDNA driven by the IL2-specific *tba-6* promoter in *shy87* mutants (**Figure 2G**). This led to a robust decrease in the number of IL2Q 2° and 3° dendrites, suggesting that *sax-1* functions cell-autonomously in IL2Q to mediate dendrite pruning (**Figure 2I-J**). NDR kinases are phosphorylated at conserved serine and threonine residues, residing in the hydrophobic motif (S279 in *C. elegans*) and the T-loop, which are essential for kinase activity (Millward, Hess, and Hemmings 1999; Tamaskovic et al. 2003). To test whether SAX-1 function requires its kinase activity, we next expressed SAX-1 cDNA with a S279A mutation (**Figure 2H**), which suppresses NDR kinase activity *in vitro* (Millward, Hess, and Hemmings 1999; Tamaskovic et al. 2003). SAX-1(S279A) expression did not rescue dendrite remodeling defects in *sax-1(shy87)* mutants in multiple transgenic lines tested (**Figure 2I-J**). Together, these data indicate that SAX-1/NDR functions cell-autonomously in IL2Q in a kinase-activity-dependent manner.

### MOB-2 and SAX-2/Furry function with SAX-1/NDR to direct IL2Q dendrite elimination

To identify proteins that function with SAX-1 in IL2Q dendrite elimination, we conducted a candidate genetic screen (**Table 1**). From this screen, we found that the conserved NDR activator MOB-2 and the binding partner SAX-2/Furry were required for the elimination of IL2Q branches. Similar to *sax-1* mutants (**Figure 3B**), *sax-2* (**Figure 3C**) and *mob-2* (**Figure 3E**) mutants exhibited selective retention of secondary and tertiary branches in adults, while quaternary branches were efficiently eliminated (**Figure 3G-I**). <u>M</u>ps*1*-<u>b</u>inder-related (MOB) proteins bind the N-terminal Regulatory (NTR) region of NDR kinases and promote their activation (Devroe et al. 2004; Bichsel et al. 2004; Hergovich 2011; Gógl et al. 2015). SAX-2/Furry is a ∼300 kD protein containing Armadillo repeats that may act as a scaffold for SAX-1/NDR (Cong et al. 2001; Chiba et al. 2009; Nagai and Mizuno 2014) and functions with NDR kinases in neurite growth and dendrite tiling (Zallen et al. 2000; Emoto et al. 2004; Gallegos and Bargmann 2004). To assess if *sax-1*, *mob-2*, and *sax-2* function in the same genetic pathway, we generated *sax-1;mob-*2 and *sax-1;sax*-2 double-mutants and compared their phenotypes to those of single mutants and controls (**Figure 3D; 3F-I**). Double-mutants did not show an additive effect, suggesting that they act within the same genetic pathway. These results suggest that a conserved SAX-1/SAX-2/MOB-2 complex is required for IL2Q dendrite pruning.

**Table 1.** Candidate genetic screen for SAX-1/NDR kinase interactors and substrates.

| Gene | Allele(s) | Mammalian Ortholog | Phenotype | References |
| --- | --- | --- | --- | --- |
| **Cell Growth and Size Regulators** |  |  |  |  |
| *sax-2* | *ky216* | Furry | IL2Q pruning defect post dauer | (He et al. 2005; Fang et al. 2010; Nagai and Mizuno 2014; Cervino et al. 2021; Roşianu et al. 2023) |
| *F09A5.4/mob-2* | *ok3273* | Mob2 | IL2Q pruning defect post dauer | (Devroe et al. 2004) |
| *yap-1* | *tm1416* | YAP1 | No phenotype observed | (L. Zhang et al. 2015; Irie, Nagai, and Mizuno 2020) |
| *rict-1* | *mg360* | Rictor | No phenotype observed | (Koike-Kumagai et al. 2009; Wu et al. 2013) |
| *unc-82* | *e1323* | NUAK1 | No phenotype observed | (Suzuki, Ogura, and Esumi 2006) |
| **Actin Regulators** |  |  |  |  |
| *unc-43* | *e408* | CaMKII | No phenotype observed | (Zallen et al. 2000; Rehberg et al. 2014) |
| *wsp-1* | *gm324* | WASP | No phenotype observed | (Natarajan et al. 2015) |
| **Endocytosis and Membrane Trafficking** |  |  |  |  |
| *sel-5* | *ok363* | Aak1 | No phenotype observed | (Ultanir et al. 2012; Roşianu et al. 2023) |
| *pifk-1* | *tm2348* | PI4KB | No phenotype observed | (Roşianu et al. 2023) |
| *mig-10* | *ok2499* | Raph1 | No phenotype observed | (Roşianu et al. 2023) |
| *F58G6.1/amph-1* | *tm1060* | BIN1 | No phenotype observed | (Roşianu et al. 2023) |
| *F54C9.11/rabi-1* | *tm2518* | Rabin8 | IL2Q pruning defect post dauer | (Ultanir et al. 2012; Chiba et al. 2013; Deretic et al. 2023; Fresquez et al. 2025) |
| *rab-11.1* | *tm2251* | Rab11A | No phenotype observed | (Chiba et al. 2013; Roşianu et al. 2023; Burguete, Song, and Ghabrial 2024) |
| *rab-11.2* | *tm2081* | Rab11A | IL2Q pruning defect post dauer | (Chiba et al. 2013; Roşianu et al. 2023; Burguete, Song, and Ghabrial 2024) |
| *rab-10* | *ok1494* | Rab10 | IL2Q branching defect at dauer | (Chiba et al. 2013; Homma and Fukuda 2016; Fresquez et al. 2025) |
| *rab-8* | *tm2991* | Rab8B | ILQ branching defect at dauer and post dauer | (Chiba et al. 2013; Homma and Fukuda 2016; Fresquez et al. 2025) |
| **Mitochondrial Quality Control** |  |  |  |  |
| *pink-1* | *ok3538* | PINK1 | No phenotype observed | (Wu et al. 2013) |

**Figure 3.**
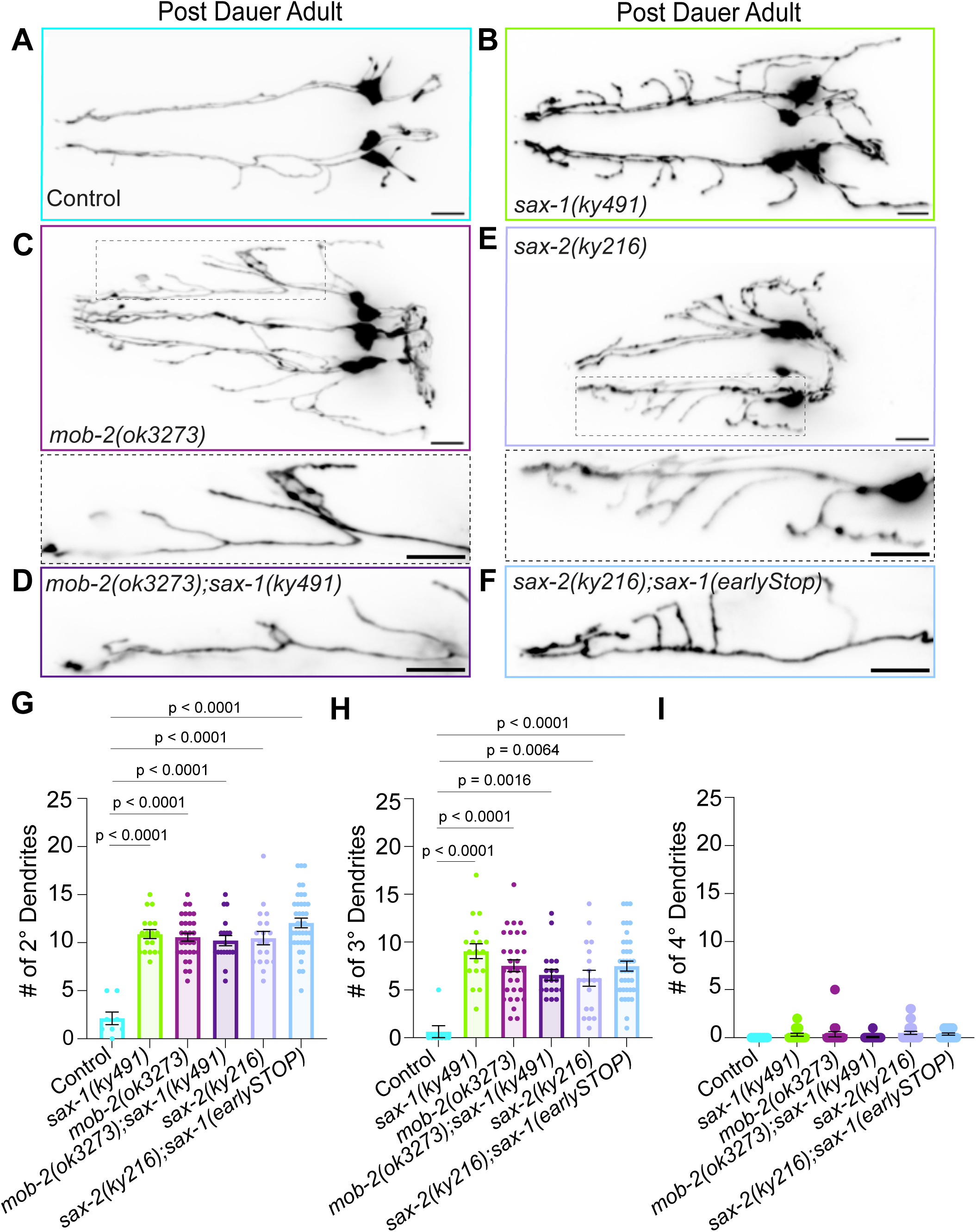
MOB-2 and SAX-2/Furry function with SAX-1/NDR to direct IL2Q dendrite elimination. Post dauer adult confocal images of (A) control and (B) *sax-1* loss-of-function, (C) *mob-2,* (D) *sax-1;mob-2* double mutants, (E) *sax-2*, and (F) *sax-1; sax-2* double-mutants. Zoomed inset of select IL2Q are shown for *mob-*2 and *sax-2* single mutants with a dashed box. Scale bars, 10μm. (G-H) Quantification of the total number of 2° (G), 3° (H), and 4° dendrites (I) in post dauer adults for genotypes shown in A-F. Brown-Forsythe and Welch ANOVA with Dunnett’s correction (2°) or Kruskal-Wallis test with Dunn’s correction (3°-4°), n = 8-39. Error bars are ±SEM, with individual data points shown.

### SAX-2/Furry localization depends on SAX-1

To gain insights into the functions of SAX-1 and SAX-2 in dendrite remodeling, we inserted three copies of spGFP11 (3xspGFP11) at the endogenous C-terminus of SAX-2 with CRISPR-Cas9 and visualized it in IL2Q by expressing spGFP1-10 under the *tba-6* promoter. Tagging of SAX-2 with spGFP11 did not lead to dendrite remodeling defects, suggesting that the tag is functional (data not shown). In control animals, SAX-2 was mostly concentrated in the cell body (data not shown), with occasional dendritic puncta observed at dauer (**Figure 4B**; **4F**). The punctate distribution of SAX-2 in IL2s closely resembles its appearance in other neurons and tissues (Gallegos and Bargmann 2004; Park et al. 2024). Interestingly, 16-20 hours after shifting dauer animals to 15°C, there was an increase in SAX-2 puncta in 1° dendrites (**Figure 4D; 4F)**, suggesting that SAX-2 may function locally during dendrite pruning. In *sax-1* mutants, we observed a robust increase of dendritic SAX-2 puncta (**Figure 4E**; **4G-H**), mostly around the 2° and 3° branchpoints, which would have been eliminated 16-20 hours following dauer recovery in control animals. This phenotype is underestimated in the quantifications (**Figure 4F-H**), in which we only measured SAX-2 puncta in the 1° dendrite, to avoid comparing dendritic arbors of different sizes. These results indicate that *sax-1* is required for the localization of SAX-2. The localization of SAX-2 in *sax-1* mutants is consistent with a failure to progress beyond an intermediate step in the branch elimination process.

**Figure 4.**
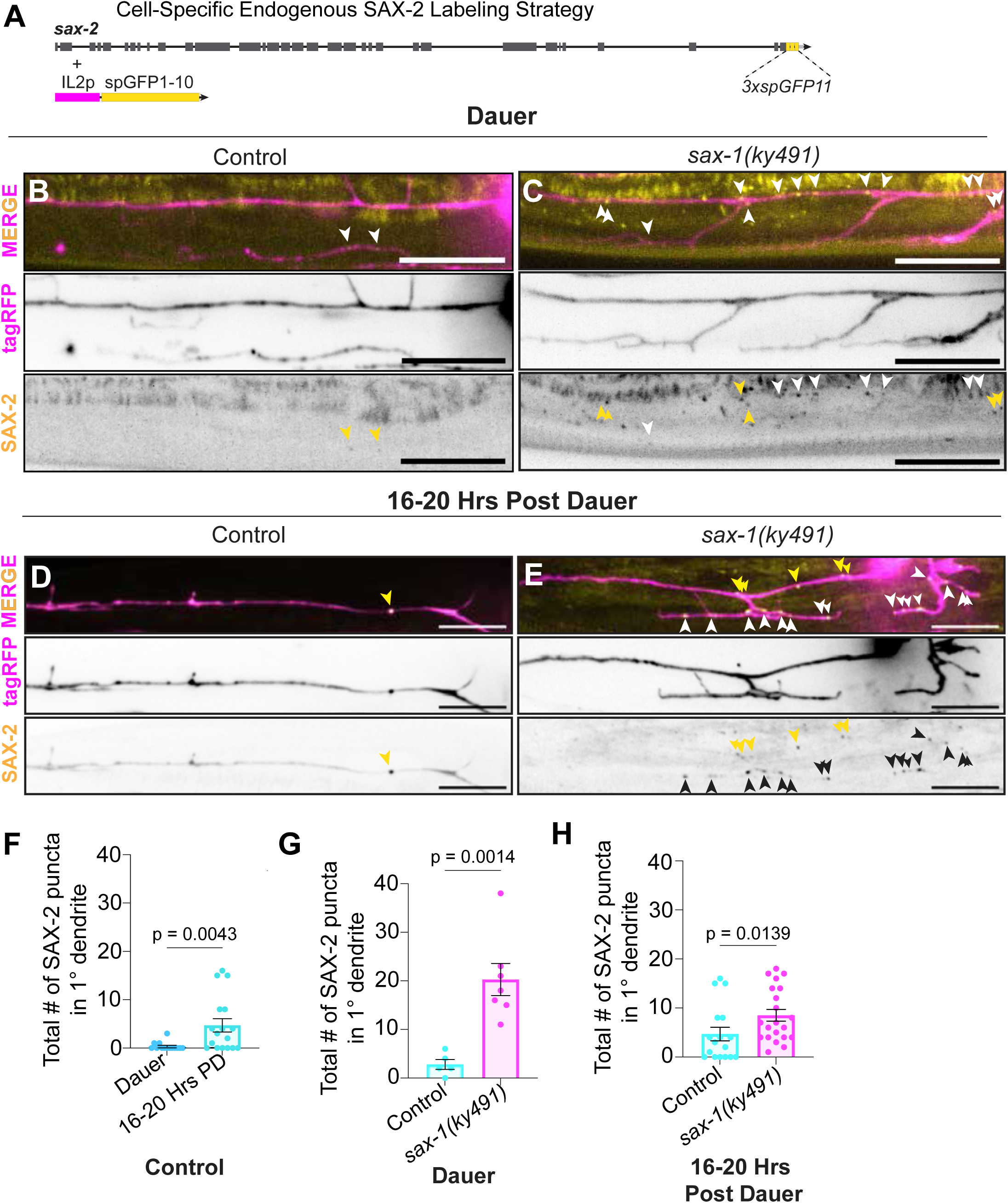
SAX-2/Furry localization depends on SAX-1 kinase. (A) Schematic illustrating the strategy to endogenously label SAX-2 in IL2 neurons using split-GFP, with GFP1-10 expressed under IL2 specific promoter. (B-C) Representative images of endogenous SAX-2::GFP (pseudo-colored yellow) in IL2 neurons (pseudo-colored magenta) at dauer arrest in control (B) and *sax-1* mutants (C). Scale bar, 10μm. (D-E) Representative IL2Q inset maximum intensity Z-projection of endogenous SAX-2::GFP in IL2 neurons 16-20 hours post dauer in control (D) and *sax-1* mutants (E). Zoomed insets show a single IL2D 1° dendrite. Yellow arrowheads denote SAX-2::GFP puncta localized at 1° dendrites; white/black arrowheads point to SAX-2::GFP puncta localized to higher-order dendrites. Scale bar, 10μm. (F) Quantification of SAX-2::GFP puncta at 1° dendrites in control animals at dauer and 16-20 hours post dauer. Mann-Whitney U test, n = 17-20. Dendritic SAX-2 puncta increase 16-20 hours following dauer recovery. (G) Quantifications of total # of SAX-2 puncta at 1° dendrites in control versus *sax-1* mutants at dauer. Unpaired *t*-test with Welch’s correction, n = 5-7. (H) Quantifications of total # of SAX-2 puncta at 1° dendrites in control versus *sax-1* mutants at 16-20 hours post dauer. Mann-Whitney U test, n = 17-21. Error bars are ±SEM.

### SAX-1 coordinates IL2Q remodeling with RABI-1/Rabin8 and RAB-11.2

Since SAX-1/NDR kinases are emerging as key regulators of membrane dynamics (Roşianu et al. 2023; Fresquez et al. 2025), we tested whether specific Rab GTPases involved in membrane recycling, along with other regulators of membrane dynamics, act as effectors of SAX-1/NDR1/2-mediated dendrite remodeling (**Table 1**). We found that mutants of RABI-1/Rabin8, a Rab8 <u>g</u>uanine-nucleotide <u>e</u>xchange factor (GEF), that was previously identified as a direct phosphorylation target of NDR kinases (Ultanir et al. 2012; Chiba et al. 2013; Roşianu et al. 2023), failed to eliminate their 2° dendrites post dauer (**Figure 5E; 5J**). Tertiary dendrites were affected (**Figure 5K**), but the phenotype was very subtle when compared to *sax-1, sax-2* or *mob-2* mutants, while 4° were efficiently eliminated (data not shown). Double-mutants of *sax-1* and *rabi-1* did not show an additive effect and phenocopied the severity of *sax-1* single mutants (**Figure 5H; 5L**). These results suggest that *rabi-1* and *sax-1* function in the same genetic pathway that coordinates secondary dendrite elimination, while *sax-1* likely directs elimination of tertiary dendrites with additional downstream regulators.

**Figure 5.**
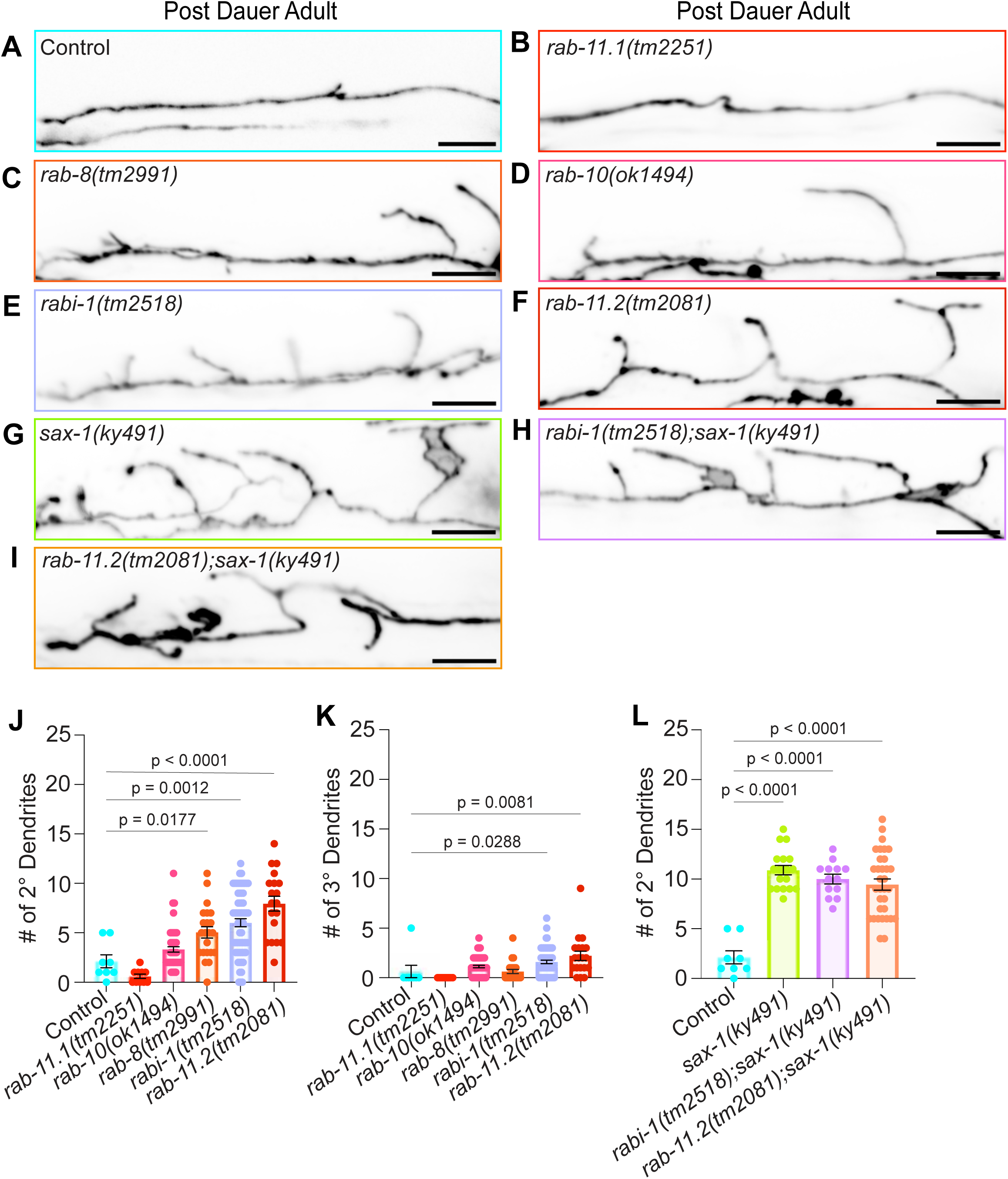
SAX-1 coordinates IL2Q remodeling with RABI-1/Rabin8 and RAB-11.2. (A-I) Representative confocal images of select IL2Q insets for control (A), *rab-11.1* (B), *rab-8* (C), *rab-10* (D), *rabi-1* (E), *rab-11.2* (F), *sax-1* (G), *rabi-1;sax-1* (H), and *rab-11.2*;*sax-1* (I) post dauer adult mutants. Scale bar, 10μm. (J-K) Quantification of total number of 2° (J) and 3° (K) dendrites for A-F. Brown-Forsythe and Welch ANOVA with Dunnett’s correction (J) or Kruskal-Wallis with Dunn’s correction (K), n = 8-62. (L) Quantification of total number of 2° dendrites in indicated genotypes. Brown-Forsythe and Welch ANOVA with Dunnett’s correction, n = 8-33. Error bars are ±SEM, with individual data points shown.

Rabin8 functions as a GEF for Rab8 and Rab10, promoting their activation during membrane trafficking (Hattula et al. 2002; Knödler et al. 2010; Feng et al. 2012; Homma and Fukuda 2016). We found that *rab-8* and *rab-10* mutants showed mild pruning defects of secondary dendrites, reminiscent to *rabi-1* mutants (**Figure 5C-D**;**5J-K**). These results suggest that RABI-1/Rabin8 likely functions with RAB-8 and RAB-10 during dendrite pruning. However, *rab-10* and *rab-8* mutants also show reduced branching at dauer (data not shown), consistent with their roles in PVD dendrite arborization (Taylor et al. 2015; Zou et al. 2015), which may influence their pruning phenotypes.

Rabin8 recruitment to recycling endosomes depends on its interaction with GTP-bound Rab11 (Chiba et al. 2013; Feng et al. 2015; Homma and Fukuda 2016; Fresquez et al. 2025). To test the involvement of Rab11 in IL2Q dendrite remodeling post dauer, we tested mutant alleles of two closely related Rab11 homologues, *rab-11.1* and *rab-11.2. rab-11.1* mutants showed grossly normal IL2Q morphology during dauer and had no pruning defect following dauer exit (**Figure 5B**; **5J-K**). In contrast, *rab-11.2* mutants showed a pruning defect following dauer exit that was similar to *rabi-1* mutants (**Figure 5F**; **5J-K**). In addition, *rab-11.2;sax-1* double mutants did not enhance the *sax-1* mutant phenotype (**Figure 5I;5L**), suggesting that they function in the same genetic pathway that eliminates secondary dendrites. Together, these results suggest that RABI-1/Rabin8 and RAB-11.2 function with SAX-1/NDR kinase to eliminate secondary dendrites, likely by coordinating membrane dynamics during IL2Q pruning.

### SAX-1 coordinates membrane retrieval

The involvement of *rab-11.2* in *sax-1-*mediated dendrite remodeling suggests that SAX-1 may function to coordinate membrane retrieval during branch elimination. To visualize endocytosis during IL2Q remodeling, we used an established genetically encoded reporter (Richardson, Yee, and Shen 2019). We expressed secreted GFP harboring a signal peptide from muscle, and expressed in IL2 neurons a chimera consisting of RFP and the transmembrane domain of mCD8 fused to an anti-GFP nanobody (GBP) (**Figure 6A**). GFP binding to GBP leads to GFP accumulating on the surface of IL2Q, and their endocytosis appears as GFP and RFP co-localized puncta (**Figure 6C-D**) (Richardson, Yee, and Shen 2019). We used a chimera lacking GBP as a control (**Figure B**) and found that this prevented the recruitment of muscle-secreted GFP to IL2Q surface or endocytic puncta (**Figure 6E-F**).

**Figure 6.**
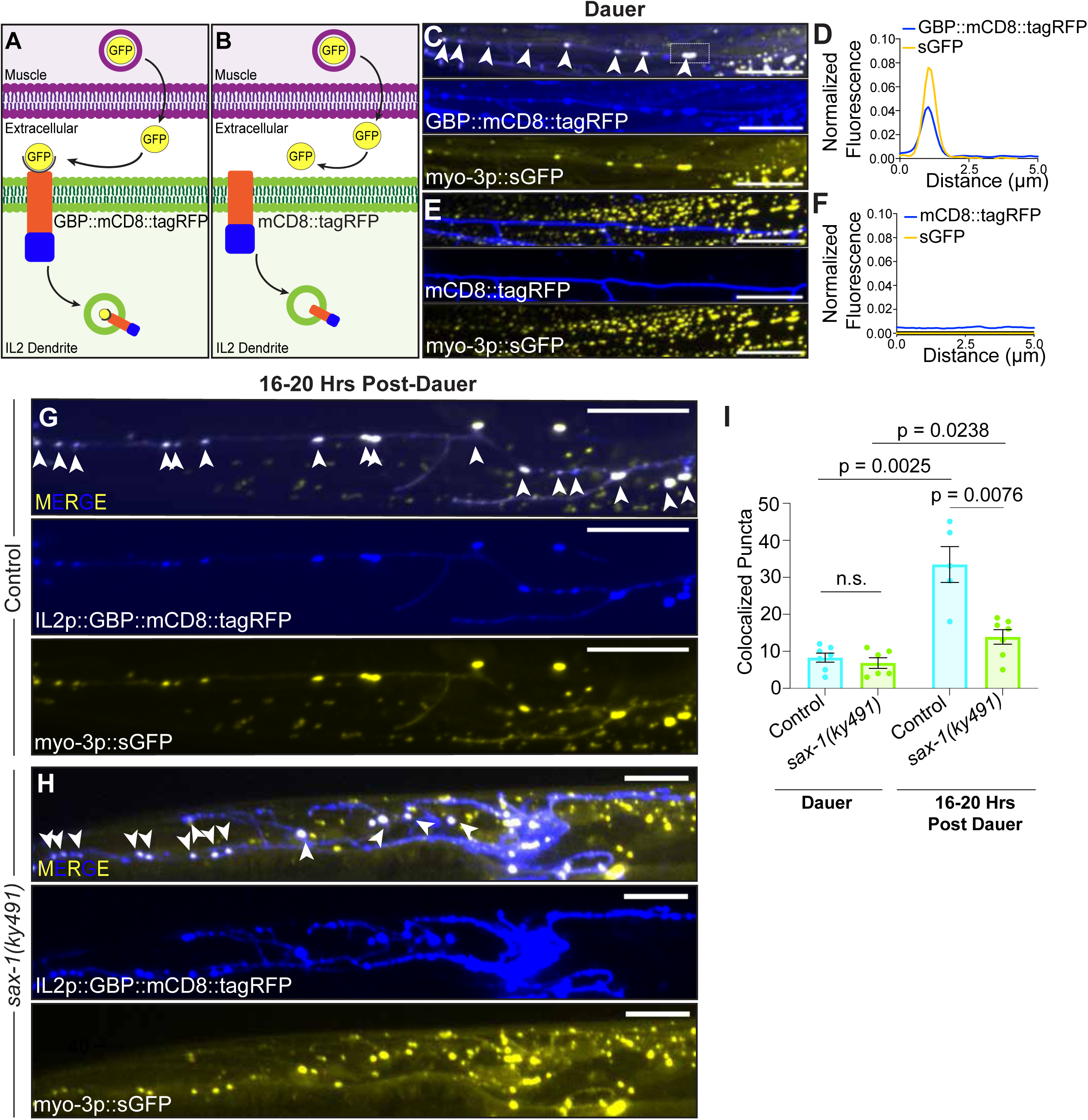
*SAX-1/NDR* promotes endocytic events during pruning. (A) Schematic of generic endocytosis reporter. mCD8::tagRFP (blue) fused to a GFP-binding nanobody are expressed under an IL2-specific promoter. Secreted GFP (yellow) from muscle binds the nanobody, enabling visualization of endocytosed GFP-mCD8 complexes as blue/yellow colocalized puncta. (B) Schematic of a control endocytosis reporter chimera lacking the GFP-binding nanobody. (C) Zoomed insets of confocal image depicting a single IL2Q 1° dendrite in control dauer with the endocytosis reporter (C-D) and a control construct lacking GFP-binding nanobody (E-F). White arrowheads show blue-yellow colocalization, suggestive of endocytosed complexes. Scale bar, 10μm. (D) Normalized fluorescence linescan of endocytic reporter showing co-localization of blue-yellow puncta (white dashed box) at dauer. (F) Normalized fluorescence linescan of control construct lacking GFP-binding nanobody showing lack of co-localization of blue-yellow puncta at dauer. (G-H) Representative zoomed insets of confocal images of the endocytosis reporter in a single IL2Q neuron at 16-20 hours post dauer in control (G) and *sax-1* mutants (H). White arrowheads depict blue-yellow co-localized puncta. Scale bar, 10μm. (I) Quantification of the total number of co-localized blue-yellow puncta at dauer and 16-20 hours post dauer in control animals and *sax-1* mutants. Mann-Whitney U test, n = 5-7. Error bars are ±SEM. *sax-1* is required for the endocytic puncta post dauer.

We found that endocytosis increases in IL2Q neurons between dauer and 16-20 hours following dauer recovery (**Figure 6I**), consistent with the idea that pruning involves membrane retrieval. There was no significant difference between the number of endocytic puncta in control and in *sax-1* mutants during dauer. Conversely, at 16-20 hours post dauer *sax-1* mutants showed a decrease in the number of endocytic events (**Figure 6I**) These results suggest that *sax-1* is specifically required for the formation or stabilization of endocytic sites during IL2Q dendrite pruning following dauer recovery.

## Discussion

The mechanisms that govern neuronal remodeling during development and under stress remain poorly understood. Here, by adapting *C. elegans* IL2Q dendrites as a model for developmentally and stress-mediated dendrite pruning, we identified a novel role for the conserved SAX-1/NDR kinase in dendrite branch-specific elimination. Our results indicate that SAX-1/NDR and its conserved interactors SAX-2/Furry and MOB-2 are required for the elimination of IL2Q secondary and tertiary, but not quaternary, branches. SAX-1/NDR is required for dendrite pruning following recovery from developmental diapause induced by manipulating *daf-7/TGF-β* or *daf-2/Insulin-receptor* signaling but is dispensable when diapause is induced by starvation. Additionally, we find that SAX-1 functions with the guanine-nucleotide exchange factor RABI-1/Rabin8 and the small GTPase RAB-11.2 to direct the elimination of 2° dendrite branches in post dauer animals. RABI-1/Rabin8 may activate small GTPases RAB-8 and RAB-10 to mediate 2° dendrite elimination and, consequently, regulate membrane retrieval. The known involvement of these proteins in membrane trafficking (Westlake et al. 2011; Ultanir et al. 2012; Chiba et al. 2013; Homma and Fukuda 2016; Fresquez et al. 2025) and our results with a genetically encoded endocytosis reporter suggest that SAX-1 promotes dendrite elimination by regulating membrane dynamics. Together, these results reveal unexpected state- and branch-specific dendrite elimination mechanisms during neuronal remodeling, governed by a conserved regulator of polarized cell growth.

SAX-1/NDR kinases are conserved regulators of polarized cell growth (Verde, Wiley, and Nurse 1998; Tamaskovic, Bichsel, and Hemmings 2003; Das et al. 2009; Gupta and McCollum 2011; Chen et al. 2019). Loss of NDR activity leads to increased cell growth in fission yeast and mammals (Hergovich et al. 2006; Das et al. 2009; Demiray et al. 2018). In *C. elegans* and *Drosophila* neurons, mutants in *sax-1* or its homolog *Trc,* respectively, fail to terminate axon growth or display excessive dendrite branching (Gallegos and Bargmann 2004; Emoto et al. 2004; Chung et al. 2016). More recently, it was shown that in hippocampal CA1 neurons, combined loss of NDR1 and NDR2 results in ectopic membrane protrusions (Roşianu et al. 2023), a phenotype that is notably similar to that of *sax-2* mutants in *C. elegans* (Park et al. 2024). Together, these results suggest a general role in restricting cell size. However, the role we identified for *sax-1* and *sax-2* in IL2Q is not in restricting cellular growth, as *sax-1* and *sax-2* mutants did not exhibit IL2Q branching defects at dauer arrest. Instead, following the temporal progression of IL2Q pruning, we found that the aberrant dendritic branches observed in mutant adults are not ectopic outgrowths, but rather the result of failed dendrite elimination. Therefore, NDR kinases are able to both inhibit growth and promote the elimination of cellular processes. How these two functions are coordinated remains to be determined. NDR kinases likely engage distinct substrates in a context-dependent manner, enabling it to differentially regulate cellular outcomes. Alternatively, since the elaboration of cellular processes such as dendritic branches involves coordinated growth and retraction (Smith et al. 2010; Shi et al. 2024), it is possible that some of the excessive branching observed in NDR kinase mutants reflects a failure in branch retraction.

How does SAX-1 promote 2° and 3° dendrite pruning? Our findings support that SAX-1 acts as a kinase and functions cell autonomously with its conserved interactors SAX-2/Furry and MOB-2. NDR kinases are activated through binding to MOB family proteins, which enhance NDR phosphorylation and stabilize their active conformation (Bichsel et al. 2004; Hergovich and and Hemmings 2005), while Furry proteins may serve as scaffolds to support kinase function (Chiba et al. 2009; Nagai and Mizuno 2014). NDR kinases can direct growth by regulating the cytoskeleton (Das et al. 2009; Norkett et al. 2020) and membrane dynamics, including regulation of exocytosis, endocytosis, membrane asymmetry, intracellular trafficking, and autophagy (Joffre et al. 2015; Yd et al. 2019; Ogura et al. 2023; Roşianu et al. 2023). We found that *sax-1* acts with *rab-11.2, rabi-1/Rabin8* and potentially *rab-8* and *rab-10* to eliminate IL2Q 2° dendrites post dauer, consistent with a role for SAX-1 in regulating membrane dynamics. The preferential elimination of 2° dendrites in these mutants suggests that membrane trafficking is differentially regulated across branch orders during pruning. We also found that *sax-1* is required for SAX-2 distribution, with an increased abundance of SAX-2 in *sax-1* mutant dendrites. If and how this localization relates to the regulation of membrane trafficking by SAX-1 is unclear. We hypothesize that the increased dendritic localization of SAX-2 is due its retention at an intermediate step in membrane retrieval. Since the compartment to which SAX-2 localizes is unclear (Park et al. 2024), future work will be required to test this hypothesis.

A key and unexpected finding of this study is that different dendritic branches rely on distinct genetic requirements for their elimination, likely reflecting the partial nature of pruning in this system. Unlike in *Drosophila* da and mushroom body γ neurons, where entire dendritic arbors are eliminated, *C. elegans* IL2Q neurons selectively eliminate only dauer-induced branches. This points to mechanisms that discriminate between primary and higher-order dendrites during pruning. Moreover, the observation that the mode of dauer induction alters the pruning mechanism underscores the complexity of this process. Leveraging this system in future work could reveal how physiological states or external cues drive branch-specific elimination programs.

## Methods

### *C. elegans* strains and maintenance

All *C. elegans* strains were cultured on <u>N</u>ematode <u>G</u>rowth <u>M</u>edium (NGM) seeded with *Escherichia coli* OP50. Animals were examined at multiple developmental stages, including normal development adults, dauer larva, 16-20hrs post dauer larva, and post dauer adults. Normal development adults were grown at 15°C. Dauer arrest was induced and dauer larva were maintained at 25°C. Post dauer animals were transferred to and maintained at 15°C, unless otherwise indicated. A detailed list of *C. elegans* strains used in the study is provided in the Key Resources Table.

### Molecular Cloning

Plasmids were constructed using conventional restriction-ligation methods or Gibson Assembly. Genes of interest were ordered as gene fragments from IDT and PCR cloned. Mutagenesis was performed using the QuikChange Lightning Multi Site-Directed Mutagenesis Kit (Agilent, Cat# 210513). Plasmids were confirmed by Sanger Sequencing. A list of plasmids used in this study are listed in the Key Resources Table. pCER229 was a gift from Claire Richardson. Plasmids were injected at the following concentrations to generate transgenic *C. elegans* lines: pRF4 (/μL), pGLS8 (15 ng/μL), pGLS9 (10ng/μL), pOVG6 (15ng/μL), pPFD12 (40ng/μL), pPFD21 (10ng/μL), pPFD37 (/μL), pPFD57 (5ng/μL), pCER229 (10ng/μL), and pPFD74 (5ng/μL). Plasmid sequences and constructs are available upon request.

### *C. elegans* transgenic strain generation

Transgenic strains were generated by microinjecting plasmid DNA into the gonads of young adult animals following established protocols (Mello and Fire 1995). Plasmids containing *unc-122p::eGFP, unc-122p::tagRFP, elt-7p::eGFP::NLS*, and *elt-7p::TagRFP-T::NLS* were used as co-injection markers. Following microinjection, F1 progeny transmitting the fluorescent reporter or co-injection marker were individually isolated, and their F2 progeny were screened under a compound microscope to identify appropriate expression levels. Integrated strains were generated using the TMP/UV method, outcrossed a minimum of six times, and subsequently mapped to specific chromosomes using standard mapping strains prior to experimental use.

### Dauer Induction and Exit

Dauer induction and exit were controlled using the temperature-sensitive, constitutive *daf-7/TGFβ* mutant allele *e1372*. Larva 4 (L4) stage animals were maintained at 25°C to allow dauer induction. Then, dauer larva were manually transferred to NGM plates with freshly seeded *E. coli* OP50 and shifted to 15°C to prompt dauer exit towards reproductive development.

### Genetic screen to isolate IL2Q pruning regulators

To identify novel IL2Q neuron pruning regulators, we conducted an unbiased forward genetic screen (**Supplemental Figure 1C**). Animals (L4 larva) carrying the temperature-sensitive *daf-7(e1372)* mutant allele were exposed to <u>e</u>thyl <u>m</u>ethane<u>s</u>ulfonate (EMS) following previous protocols (Brenner 1974) to induce random genomic point mutations. After mutagenesis, animals were maintained at 15°C to allow for normal development. F1 animals (∼3 per plate) were then transferred to 25°C to induce dauer arrest in F2 progeny. Dauer larvae were then manually transferred to NGM plates with freshly seeded *E. coli* OP50 and incubated at 15°C to prompt dauer exit. Nematodes at the post dauer day 1 adult stage were subsequently mounted on a 2% agarose pad, immobilized with 10mM levamisole dissolved in M9, and screened under a fluorescent compound microscope for IL2Q remodeling defects. Mutant candidates were isolated and outcrossed at least once to our control strain to confirm the heritability of the observed phenotypes. The causal variant for *shy87*, described in this study, were mapped by whole-genome sequencing and SNV linkage mapping with the MiModD software package run in a Galaxy server. To further confirm the identity of the isolated shy87 allele, we employed a transgenic rescue strategy utilizing a *sax-1* containing fosmid and were able to successfully rescue the mutant phenotype.

### IL2Q Remodeling Timeseries

Synchronized populations of dauer larva induced at 25°C, carrying a temperature-sensitive *daf-7(e1372)* mutation, were manually transferred to NGM plates with freshly seeded *E. coli* OP50 and incubated at 15°C to prompt dauer exit towards reproductive development. At 8hrs post-transfer, we screened under a dissecting microscope and selected for larva that exhibited pharyngeal pumping and foraging behavior. The larvae were then kept at 15°C for further examination. Animals were then mounted on 2% agarose pads and 10mM levamisole dissolved in M9 for imaging at defined timepoints (12, 14, 16, 18, 20, 22, 24, 26, 28, 30 hours). Image acquisition settings were maintained throughout all timepoints. Quantification of remodeling events was performed as described in the IL2Q Neurite Scoring section.

### Fluorescence microscopy and sample preparation

Nematodes were mounted on a 2% agarose pad and paralyzed in a droplet of 10mM Levamisole dissolved in M9 buffer. Animals were imaged at the following developmental stages: adults (normal development), dauer larva, 16-20hrs post dauer larva, and post dauer old adults, as indicated in the Figures. Images were acquired with a Laser Sage DMi8 inverted microscope (Leica) equipped with a VT-iSIM system (BioVision) and an ORCA-Flash 4.0 camera (Hamamatsu) controlled by MetaMorph Advanced Confocal Acquisition Software Package. The microscope was equipped with an HC PL APO 100x/1.47NA Oil, HC PL APO 63x/1.40NA Oil CS2, a HC PL APO 40x/1.30NA Oil CS2, and a HC PL APO 20x/0.8NA Air objective. Maximum intensity projections were generated in FIJI (ImageJ2).

### CRISPR/Cas9 Genome Editing

All CRISPR-Cas9-engineered strains were generated following the Mello Lab published protocol (Dokshin et al. 2018). We synthesized sgRNA from single-stranded DNA utilizing the EnGen sgRNA Synthesis Kit, *S. pygenes* (NEB Cat#E3322S), followed by sgRNA purification utilizing the Monarch RNA cleanup kit (NEB #T2040). A detailed list of sgRNA target sites is provided in the Key Resources Table, along with corresponding repair templates listed.

### IL2Q Neurite Scoring

Raw imaging files were imported to ImageJ2 for manual IL2Q neurite scoring. Scoring was based on the stereotypical arborization pattern of IL2Q neurons (**Supplemental Figure 1A**). In control animals, primary (1°) dendrites extend anteriorly from the cell body toward the nose along the pharyngeal axis. Secondary (2°) dendrites emerge perpendicularly from the 1° dendrite, extending toward the ventral and dorsal midlines. Tertiary (3°) dendrites arise from the 2° dendrites and bifurcate along the anterior-posterior axis. Quaternary (4°) dendrites extend perpendicularly from the 3° dendrites toward the body-wall muscle quadrants. In addition to anterior arborization, neurons also project posteriorly directed neurites, but these were excluded. For each animal, IL2D and IL2V pairs (four) neurons and all dendrite orders (1°–4°) were scored, and data were compiled to determine the total number of 2°, 3°, and 4° dendrites per genotype. For adults and 16-20hrs animals, the previously described scoring strategy was employed.

### Fluorescence Quantification

Following acquisition, raw imaging files were processed in ImageJ2. For quantification of endogenous and cell-specific SAX-2 puncta in control and *sax-1* mutants, we manually counted GFP(+) puncta across the length of the IL2D/V primary (1°) dendrite. For quantification of the generic endocytosis reporter, normalized fluorescence linescans were generated by tracing the length of an IL2D/V primary (1°) dendrite and measuring fluorescence intensity along the neurite using the *Plot Profile* function. Colocalized puncta between secreted GFP and IL2-expressed mCD8::tagRFP were manually counted from images of live animals.

### Statistical Analysis

Statistical analyses were performed on GraphPad Prism 10. Data were tested for normality, and parametric or nonparametric tests used as appropriate. Statistical significance was determined using unpaired *t*-test, unpaired *t*-test with Welch’s correction, or Mann-Whitney U test. For multiple comparisons, we used Brown-Forsythe and Welch ANOVA test or Kruskal-Wallis test followed by a post test. Sample sizes are described in figure legends. Error bars are represented as ± SEM, with individual data points shown.

## Key Resources Table

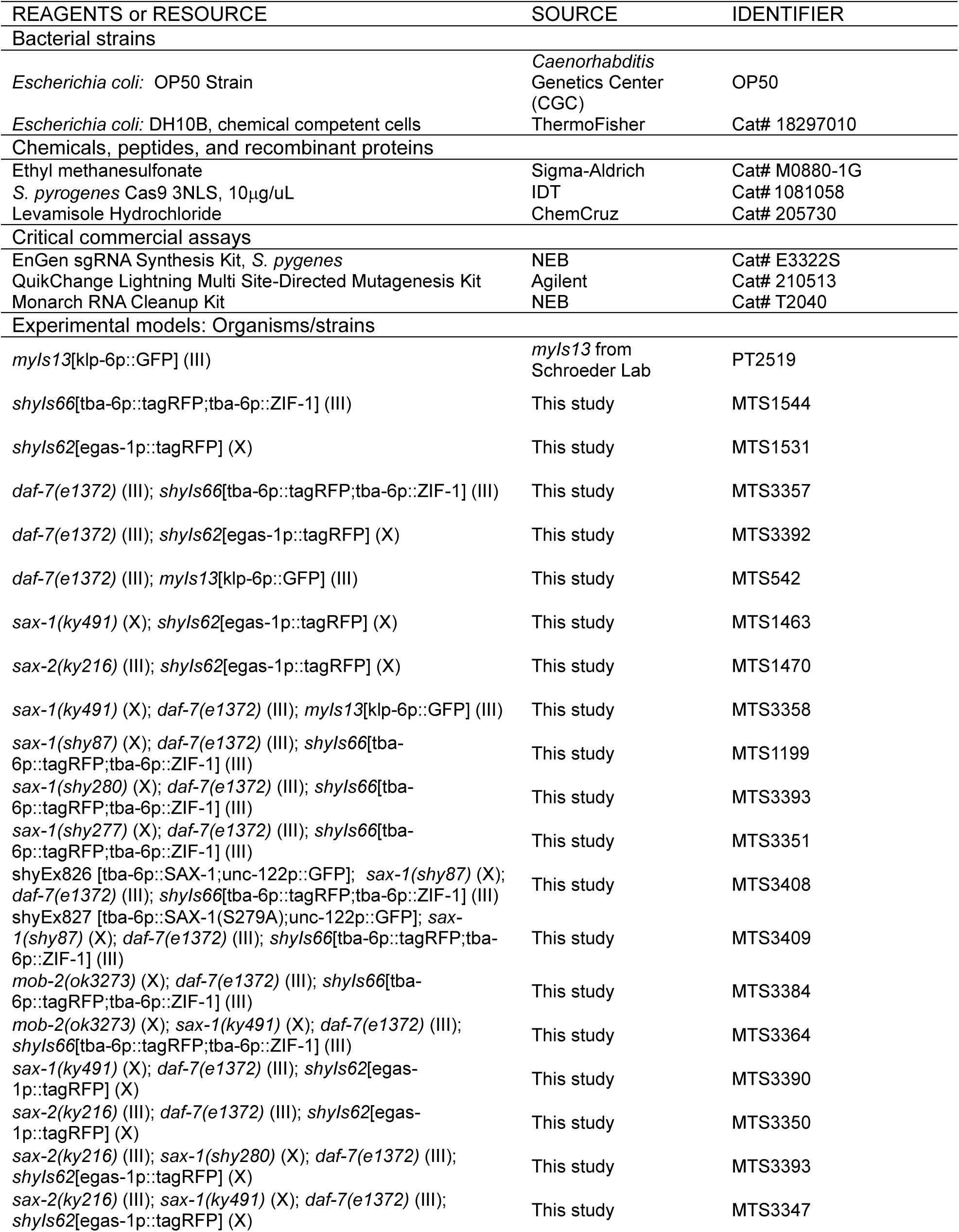
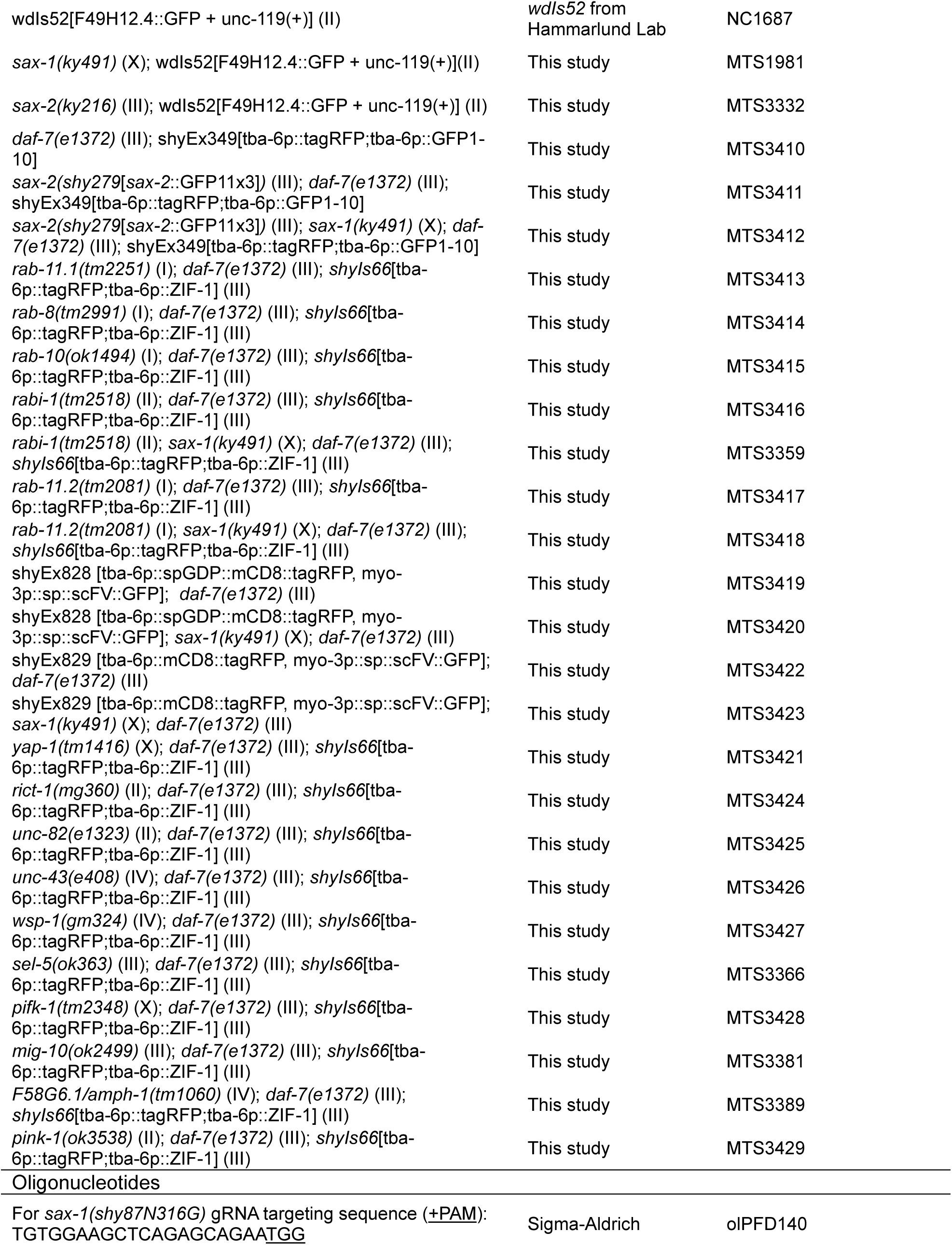
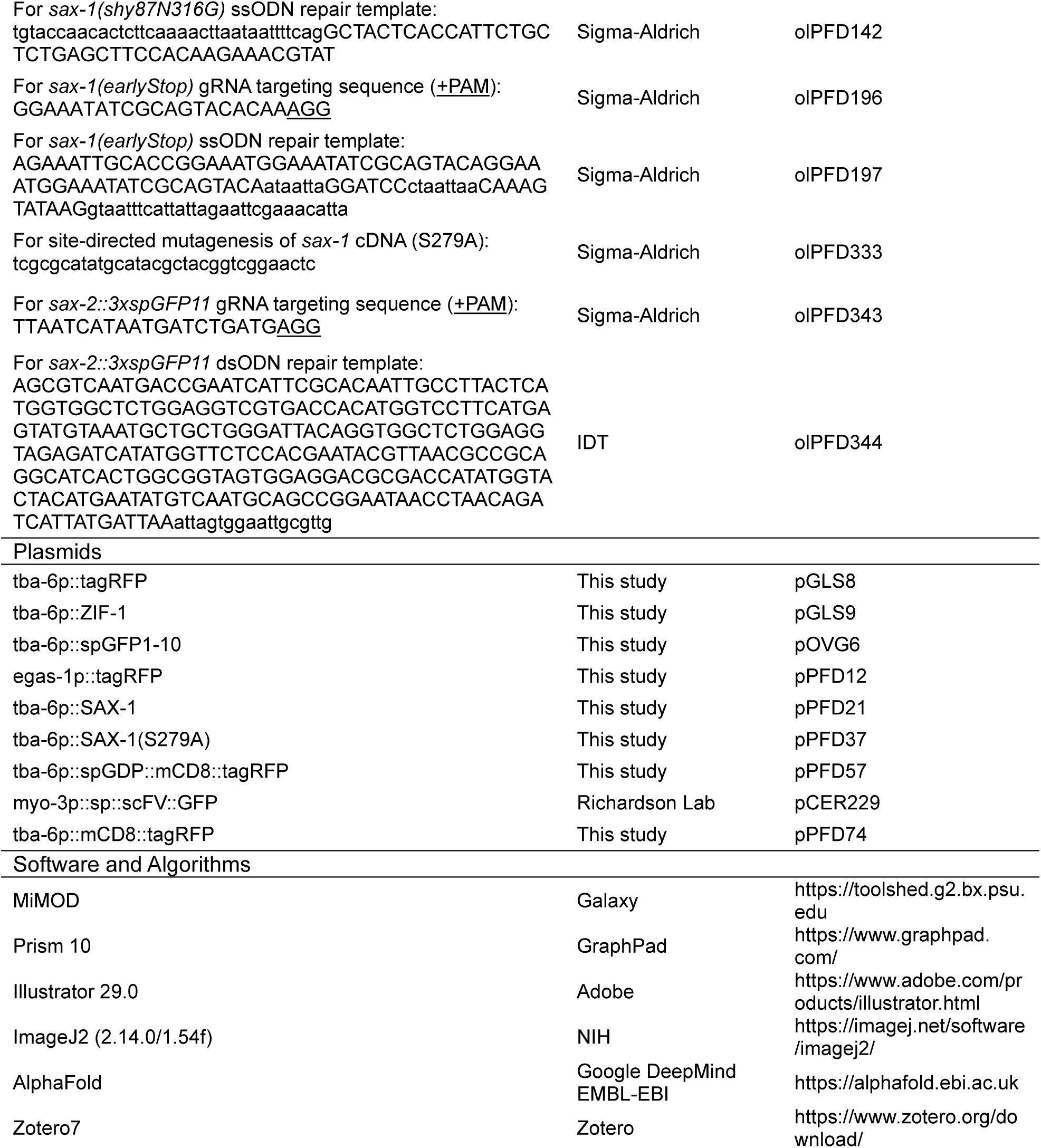

## Resource Availability

### Lead Contact

Further information and requests for reagents should be directed to the lead contact: Shaul Yogev.

## Materials Availability

Plasmids and transgenic *C. elegans* strains generated for this study are available form the lead contact upon request.

## Data and Code Availability

Raw data used for this study are available from the lead contact upon request.

## Acknowledgements

We thank members of the Yogev lab for valuable input and technical advice. We thank OVG and GLS for plasmids. We acknowledge the Yale Center for Genome Analysis, which is supported by the NIH National Institute of General Medical Science (1S10OD030363-01A1). We thank Claire Richardson (University of Wisconsin, Madison) for generously sharing the following plasmids: pCER206 and pCER229. Some strains provided by *Caenorhabditis* Genome Center (CGC), which is funded by NIH Office of Research Infrastructure Programs (P40 OD010440). We acknowledge the Mitani Lab, the National Bioresource Project for the Nematode (Japan) for alleles provided. This work was supported by R35GM133573 to SY and NIH F31-NS122294 and GE016776 to PVFD.

## Author Contributions

P.V.F.D. and S.Y. conceptualized and designed the experiments described in this study. P.V.F.D executed the experiments. P.V.F.D and S.Y. wrote, reviewed, and edited the manuscript.

## Declaration of Interests

The authors declare no competing interests.

---

**Supplemental Figure 1.**
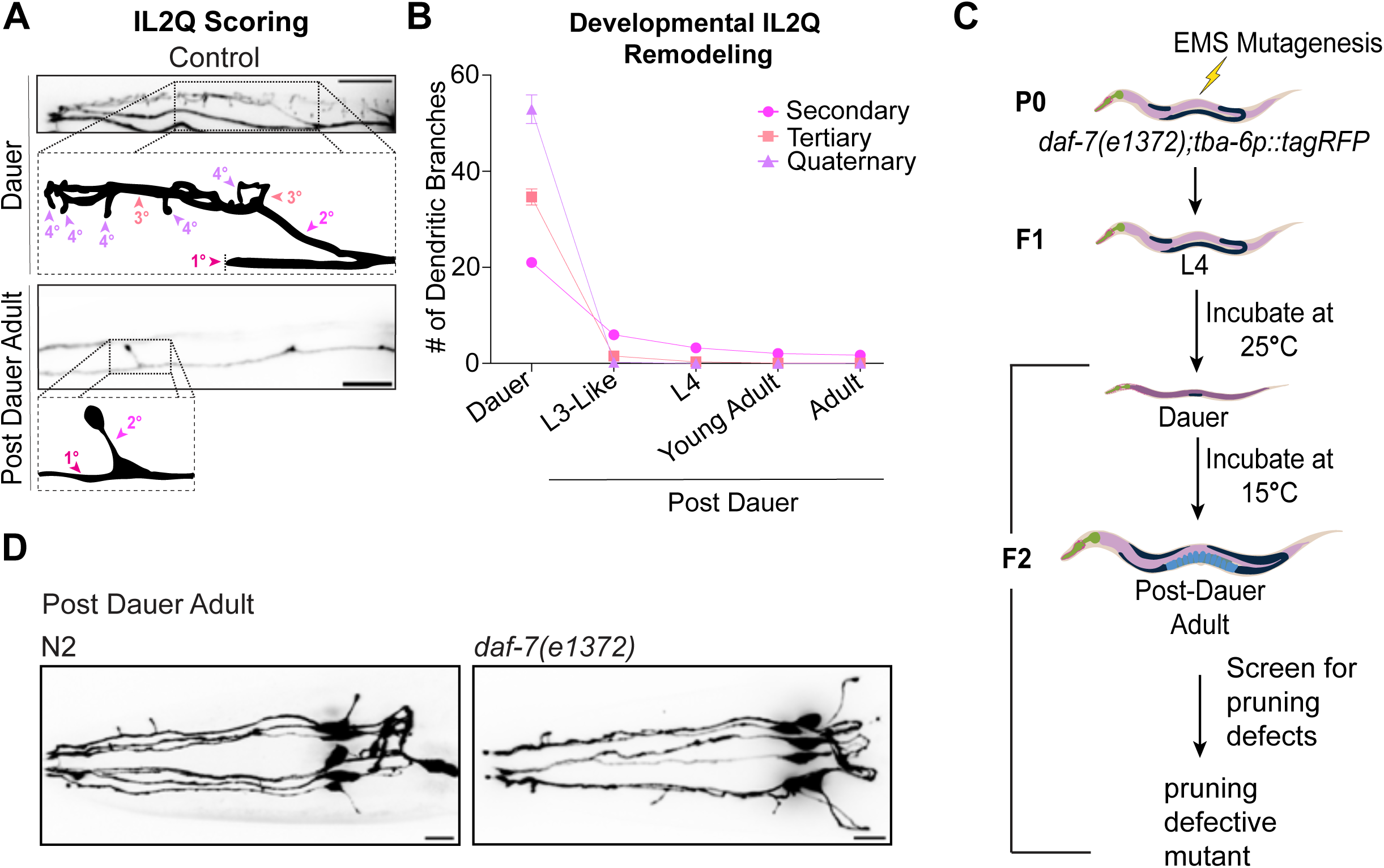
(A) IL2Q scoring: Representative Z-projection insets of select IL2Q neurons in control animals at dauer and post dauer adults, with schematic overlays indicating (with arrowhead) branch order (1°-4°) used for scoring dendrites. Scale bar, 10μm. (B) Line graph depicting the progressive elimination of IL2Q higher-order dendrites following dauer recovery towards adulthood, n = 11-53. (C) Schematic depicting strategy for the genetic screen. (D) Representative maximum intensity Z-projections of post dauer adults in N2 (left) and *daf-7(e1372)* mutants (right) show no observable difference in post dauer adults IL2Q morphology or pruning. Scale bar, 10μm.

**Supplemental Figure 2.**
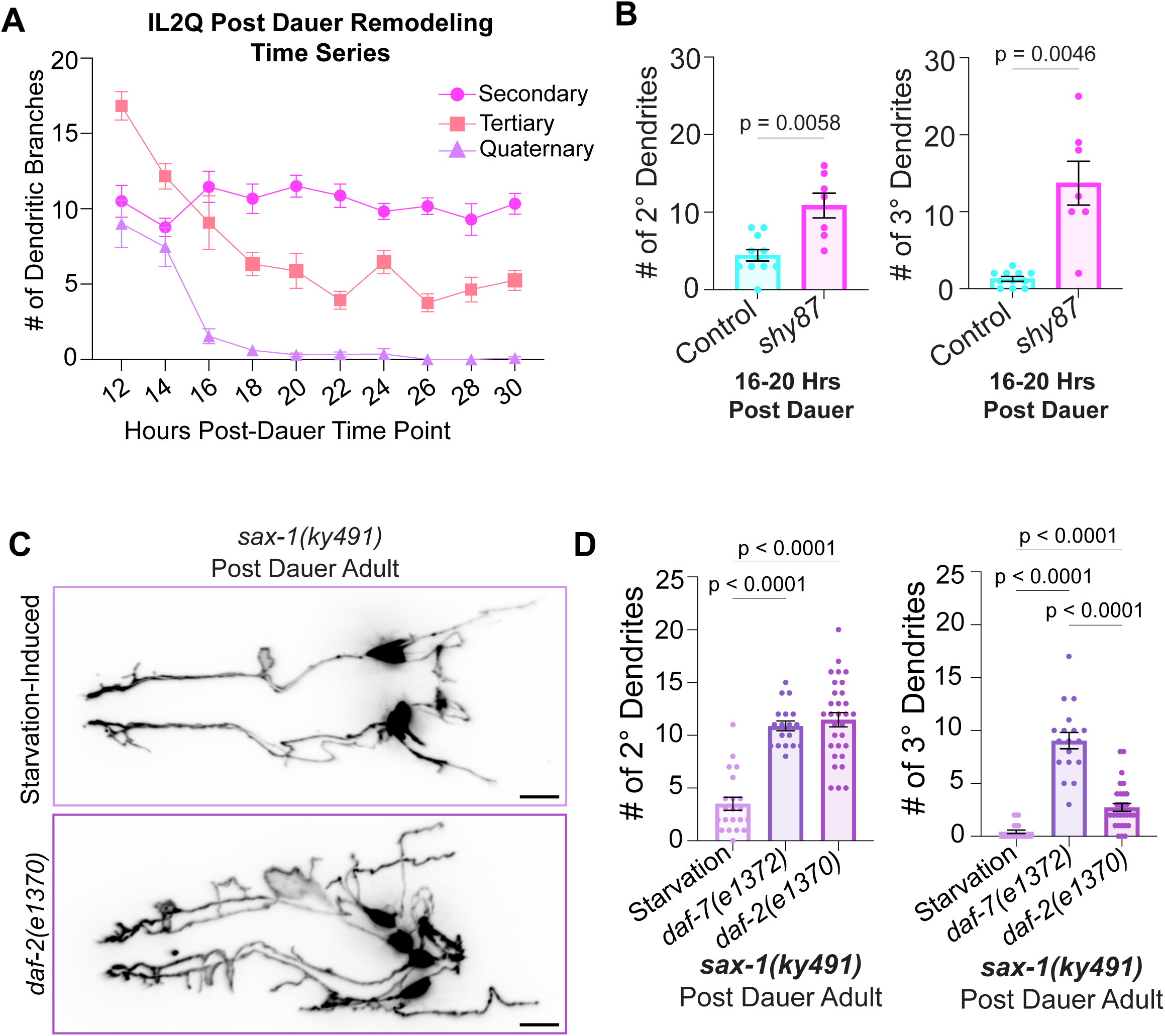
s*a*x*-1* is required for branch elimination, not inhibiting branch growth. (A) Line graph depicting the elimination of IL2Q higher-order dendrites over time (hours) post dauer (transfer to 15°C) in control animals, n = 6-17. (B) Quantification of total 2° and 3° dendrite numbers in control and *shy87* mutants at 16-20 hrs post dauer. Unpaired *t*-test with Welch’s correction, n = 7-11. (C) Representative Z-projection of *sax-1(ky491)* post dauer adults following dauer induction via starvation (top) or a dauer constitutive mutant, *daf-2/Insulin-receptor* (bottom). Scale bar, 10μm. (D) Quantification of 2° and 3° dendrites of *sax-1(ky491)* mutants following dauer induction via *daf-7(e1372)*, *daf-2(e1370)*, and starvation conditions. Brown-Forsythe and Welch ANOVA with Dunnett’s correction, n = 18-31. Error bars are ±SEM, with individual data points shown.

## Notes

### Competing Interest Statement

The authors have declared no competing interest.

## References

1. Altun, Z.F. and Hall, D.H. 2009. “Introduction.” WormAtlas. 10.3908/wormatlas.1.1

2. Androwski, Rebecca J., Kristen M. Flatt, and Nathan E. Schroeder. 2017. “Phenotypic Plasticity and Remodeling in the Stress-Induced Caenorhabditis Elegans Dauer.” Wiley Interdisciplinary Reviews. Developmental Biology 6 (5). 10.1002/wdev.278.

3. Bichsel, Samuel J., Rastislav Tamaskovic, Mario R. Stegert, and Brian A. Hemmings. 2004. “Mechanism of Activation of NDR (Nuclear Dbf2-Related) Protein Kinase by the hMOB1 Protein*.” Journal of Biological Chemistry 279 (34): 35228–35. 10.1074/jbc.M404542200.

4. Brenner, S. 1974. “THE GENETICS OF CAENORHABDITIS ELEGANS.” Genetics 77 (1): 71–94. 10.1093/genetics/77.1.71.

5. Brunson, Kristen L., Enikö Kramár, Bin Lin, Yuncai Chen, Laura Lee Colgin, Theodore K. Yanagihara, Gary Lynch, and Tallie Z. Baram. 2005. “Mechanisms of Late-Onset Cognitive Decline after Early-Life Stress.” Journal of Neuroscience 25 (41): 9328–38. 10.1523/JNEUROSCI.2281-05.2005.

6. Burguete, Alondra S., Yanjun Song, and Amin S. Ghabrial. 2024. “The Ccm3-GckIII Signaling Axis Regulates Rab11-Dependent Recycling to the Apical Compartment.” bioRxiv. 10.1101/2024.05.03.592387.

7. Burnell, Ann M., Koen Houthoofd, Karen O’Hanlon, and Jacques R. Vanfleteren. 2005. “Alternate Metabolism during the Dauer Stage of the Nematode *Caenorhabditis Elegans*.” *Experimental Gerontology*, Metabolism, Aging and Longevity, 40 (11): 850–56. 10.1016/j.exger.2005.09.006.

8. Cassada, Randall C., and Richard L. Russell. 1975. “The Dauerlarva, a Post-Embryonic Developmental Variant of the Nematode *Caenorhabditis Elegans*.” Developmental Biology 46 (2): 326–42. 10.1016/0012-1606(75)90109-8.

9. Cervino, Ailen S., Bruno Moretti, Carsten Stuckenholz, Hernán E. Grecco, Lance A. Davidson, and M. Cecilia Cirio. 2021. “Furry Is Required for Cell Movements during Gastrulation and Functionally Interacts with NDR1.” Scientific Reports 11 (1): 6607. 10.1038/s41598-021-86153-x.

10. Chen, Chuan, Marbelys Rodriguez Pino, Patrick Roman Haller, and Fulvia Verde. 2019. “Conserved NDR/LATS Kinase Controls RAS GTPase Activity to Regulate Cell Growth and Chronological Lifespan.” Molecular Biology of the Cell 30 (20): 2598–2616. 10.1091/mbc.E19-03-0172.

11. Chiba, Shuhei, Yuta Amagai, Yuta Homma, Mitsunori Fukuda, and Kensaku Mizuno. 2013. “NDR2- mediated Rabin8 Phosphorylation Is Crucial for Ciliogenesis by Switching Binding Specificity from Phosphatidylserine to Sec15.” The EMBO Journal 32 (6): 874–85. 10.1038/emboj.2013.32.

12. Chiba, Shuhei, Masanori Ikeda, Kokichi Katsunuma, Kazumasa Ohashi, and Kensaku Mizuno. 2009. “MST2- and Furry-Mediated Activation of NDR1 Kinase Is Critical for Precise Alignment of Mitotic Chromosomes.” Current Biology 19 (8): 675–81. 10.1016/j.cub.2009.02.054.

13. Christian, K. M., A. D. Miracle, C. L. Wellman, and K. Nakazawa. 2011. “Chronic Stress-Induced Hippocampal Dendritic Retraction Requires CA3 NMDA Receptors.” Neuroscience 174 (February):26–36. 10.1016/j.neuroscience.2010.11.033.

14. Chung, Samuel H., Mehraj R. Awal, James Shay, Melissa M. McLoed, Eric Mazur, and Christopher V. Gabel. 2016. “Novel DLK-Independent Neuronal Regeneration in Caenorhabditis Elegans Shares Links with Activity-Dependent Ectopic Outgrowth.” Proceedings of the National Academy of Sciences of the United States of America 113 (20): E2852–2860. 10.1073/pnas.1600564113.

15. Cong, Jingli, Wei Geng, Biao He, Jingchun Liu, Jeannette Charlton, and Paul N. Adler. 2001. “The Furry Gene of Drosophila Is Important for Maintaining the Integrity of Cellular Extensions during Morphogenesis.” Development 128 (14): 2793–2802. 10.1242/dev.128.14.2793.

16. Das, Maitreyi, David J. Wiley, Xi Chen, Kavita Shah, and Fulvia Verde. 2009. “The Conserved NDR Kinase Orb6 Controls Polarized Cell Growth by Spatial Regulation of the Small GTPase Cdc42.” Current Biology: CB 19 (15): 1314–19. 10.1016/j.cub.2009.06.057.

17. Demiray, Yunus E., Kati Rehberg, Stefanie Kliche, and Oliver Stork. 2018. “Ndr2 Kinase Controls Neurite Outgrowth and Dendritic Branching Through Α1 Integrin Expression.” Frontiers in Molecular Neuroscience 11 (March). 10.3389/fnmol.2018.00066.

18. Deretic, Dusanka, Beatrice M Tam, Orson L Moritz, Michael Robichaux, and Theresa Fresquez. 2023. “Rabin8 Phosphorylation by NDR2 Directs Membrane Progression in the Ciliary Trafficking of Rhodopsin.” Investigative Ophthalmology & Visual Science 64 (8): 4443.

19. Devroe, Eric, Hediye Erdjument-Bromage, Paul Tempst, and Pamela A. Silver. 2004. “Human Mob Proteins Regulate the NDR1 and NDR2 Serine-Threonine Kinases *.” Journal of Biological Chemistry 279 (23): 24444–51. 10.1074/jbc.M401999200.

20. Dokshin, Gregoriy A., Krishna S. Ghanta, Katherine M. Piscopo, and Craig C. Mello. 2018. “Robust Genome Editing with Short Single-Stranded and Long, Partially Single-Stranded DNA Donors in Caenorhabditis Elegans.” Genetics 210 (3): 781–87. 10.1534/genetics.118.301532.

21. Emoto, Kazuo, Ying He, Bing Ye, Wesley B. Grueber, Paul N. Adler, Lily Yeh Jan, and Yuh-Nung Jan. 2004. “Control of Dendritic Branching and Tiling by the Tricornered-Kinase/Furry Signaling Pathway in Drosophila Sensory Neurons.” Cell 119 (2): 245–56. 10.1016/j.cell.2004.09.036.

22. Fang, Xiaolan, Qiuheng Lu, Kazou Emoto, and Paul N. Adler. 2010. “The Drosophila Fry Protein Interacts with Trc and Is Highly Mobile in Vivo.” BMC Developmental Biology 10 (1): 40. 10.1186/1471-213X-10-40.

23. Feng, Shanshan, Andreas Knödler, Jinqi Ren, Jian Zhang, Xiaoyu Zhang, Yujuan Hong, Shaohui Huang, Johan Peränen, and Wei Guo. 2012. “A Rab8 Guanine Nucleotide Exchange Factor-Effector Interaction Network Regulates Primary Ciliogenesis.” The Journal of Biological Chemistry 287 (19): 15602–9. 10.1074/jbc.M111.333245.

24. Feng, Shanshan, Bin Wu, Johan Peränen, and Wei Guo. 2015. “Kinetic Activation of Rab8 Guanine Nucleotide Exchange Factor Rabin8 by Rab11.” In Rab GTPases: Methods and Protocols, edited by Guangpu Li, 99–106. New York, NY: Springer. 10.1007/978-1-4939-2569-8_8.

25. Fresquez, Theresa, Beatrice M. Tam, Shannon C. Eshelman, Orson L. Moritz, Michael A. Robichaux, and Dusanka Deretic. 2025. “Rabin8 Phosphorylated by NDR2, the Canine Early Retinal Degeneration Gene Product, Directs Rhodopsin Golgi-to-Cilia Trafficking.” Journal of Cell Science 138 (2): JCS263401. 10.1242/jcs.263401.

26. Furusawa, Kotaro, and Kazuo Emoto. 2021. “Spatiotemporal Regulation of Developmental Neurite Pruning: Molecular and Cellular Insights from *Drosophila* Models.” *Neuroscience Research*, Toward understanding of integral processes of circuit formation, maintenance and elimination, 167 (June):54–63. 10.1016/j.neures.2020.11.010.

27. Gallegos, Maria E., and Cornelia I. Bargmann. 2004. “Mechanosensory Neurite Termination and Tiling Depend on SAX-2 and the SAX-1 Kinase.” Neuron 44 (2): 239–49. 10.1016/j.neuron.2004.09.021.

28. Geng, Wei, Biao He, Mina Wang, and Paul N Adler. 2000. “The Tricornered Gene, Which Is Required for the Integrity of Epidermal Cell Extensions, Encodes the Drosophila Nuclear DBF2-Related Kinase.” Genetics 156 (4): 1817–28. 10.1093/genetics/156.4.1817.

29. Gerisch, Birgit, Cindy Weitzel, Corinna Kober-Eisermann, Veerle Rottiers, and Adam Antebi. 2001. “A Hormonal Signaling Pathway Influencing *C. Elegans* Metabolism, Reproductive Development, and Life Span.” Developmental Cell 1 (6): 841–51. 10.1016/S1534-5807(01)00085-5.

30. Gógl, Gergő, Kyle D. Schneider, Brian J. Yeh, Nashida Alam, Alex N. Nguyen Ba, Alan M. Moses, Csaba Hetényi, Attila Reményi, and Eric L. Weiss. 2015. “The Structure of an NDR/LATS Kinase–Mob Complex Reveals a Novel Kinase–Coactivator System and Substrate Docking Mechanism.” PLOS Biology 13 (5): e1002146. 10.1371/journal.pbio.1002146.

31. Golden, James W., and Donald L. Riddle. 1982. “A Pheromone Influences Larval Development in the Nematode Caenorhabditis Elegans.” Science 218 (4572): 578–80. 10.1126/science.6896933.

32. Golden, James W., and Donald L. Riddle.1984. “The *Caenorhabditis Elegans* Dauer Larva: Developmental Effects of Pheromone, Food, and Temperature.” Developmental Biology 102 (2): 368–78. 10.1016/0012-1606(84)90201-X.

33. Gottlieb, S, and G Ruvkun. 1994. “Daf-2, Daf-16 and Daf-23: Genetically Interacting Genes Controlling Dauer Formation in Caenorhabditis Elegans.” Genetics 137 (1): 107–20. 10.1093/genetics/137.1.107.

34. Gupta, Sneha, and Dannel McCollum. 2011. “Crosstalk between NDR Kinase Pathways Coordinates Cell Cycle Dependent Actin Rearrangements.” Cell Division 6 (November):19. 10.1186/1747-1028-6-19.

35. Hattula, Katarina, Johanna Furuhjelm, Airi Arffman, and Johan Peränen. 2002. “A Rab8-Specific GDP/GTP Exchange Factor Is Involved in Actin Remodeling and Polarized Membrane Transport.” Molecular Biology of the Cell 13 (9): 3268–80. 10.1091/mbc.e02-03-0143.

36. He, Ying, Xiaolan Fang, Kazuo Emoto, Yuh-Nung Jan, and Paul N. Adler. 2005. “The Tricornered Ser/Thr Protein Kinase Is Regulated by Phosphorylation and Interacts with Furry during Drosophila Wing Hair Development.” Molecular Biology of the Cell 16 (2): 689–700. 10.1091/mbc.E04-09-0828.

37. Hergovich, Alexander. 2011. “MOB Control: Reviewing a Conserved Family of Kinase Regulators.” Cellular Signalling 23 (9): 1433–40. 10.1016/j.cellsig.2011.04.007.

38. Hergovich, Alexander, Bichsel, Samuel J., and Brian A. and Hemmings. 2005. “Human NDR Kinases Are Rapidly Activated by MOB Proteins through Recruitment to the Plasma Membrane and Phosphorylation.” Molecular and Cellular Biology 25 (18): 8259–72. 10.1128/MCB.25.18.8259-8272.2005.

39. Hergovich, Alexander, Mario R. Stegert, Debora Schmitz, and Brian A. Hemmings. 2006. “NDR Kinases Regulate Essential Cell Processes from Yeast to Humans.” Nature Reviews Molecular Cell Biology 7 (4): 253–64. 10.1038/nrm1891.

40. Homma, Yuta, and Mitsunori Fukuda. 2016. “Rabin8 Regulates Neurite Outgrowth in Both GEF Activity-Dependent and -Independent Manners.” Molecular Biology of the Cell 27 (13): 2107–18. 10.1091/mbc.E16-02-0091.

41. Hu, Patrick J. 2018. “Dauer.” In WormBook: The Online Review of C. Elegans Biology [Internet]. WormBook. https://www.ncbi.nlm.nih.gov/books/NBK116082/.

42. Hurd, Daryl D, Renee M Miller, Lizbeth Núñez, and Douglas S Portman. 2010. “Specific α- and β-Tubulin Isotypes Optimize the Functions of Sensory Cilia in Caenorhabditis Elegans.” Genetics 185 (3): 883–96. 10.1534/genetics.110.116996.

43. Irie, K., T. Nagai, and K. Mizuno. 2020. “Furry Protein Suppresses Nuclear Localization of Yes-Associated Protein (YAP) by Activating NDR Kinase and Binding to YAP.” J Biol Chem 295 (10): 3017–28. 10.1074/jbc.RA119.010783.

44. Joffre, Carine, Nicolas Dupont, Lily Hoa, Valenti Gomez, Raul Pardo, Catarina Gonçalves-Pimentel, Pauline Achard, et al. 2015. “The Pro-Apoptotic STK38 Kinase Is a New Beclin1 Partner Positively Regulating Autophagy.” Current Biology: CB 25 (19): 2479–92. 10.1016/j.cub.2015.08.031.

45. Kanamori, Takahiro, Makoto I. Kanai, Yusuke Dairyo, Kei-ichiro Yasunaga, Rei K. Morikawa, and Kazuo Emoto. 2013. “Compartmentalized Calcium Transients Trigger Dendrite Pruning in Drosophila Sensory Neurons.” Science 340 (6139): 1475–78. 10.1126/science.1234879.

46. Kanamori, Takahiro, Jiro Yoshino, Kei-ichiro Yasunaga, Yusuke Dairyo, and Kazuo Emoto. 2015. “Local Endocytosis Triggers Dendritic Thinning and Pruning in Drosophila Sensory Neurons.” Nature Communications 6 (1): 6515. 10.1038/ncomms7515.

47. Karp, Xantha. 2018. “Working with Dauer Larvae.” In WormBook: The Online Review of C. Elegans Biology [Internet]. WormBook. https://www.ncbi.nlm.nih.gov/books/NBK535516/.

48. Knödler, Andreas, Shanshan Feng, Jian Zhang, Xiaoyu Zhang, Amlan Das, Johan Peränen, and Wei Guo. 2010. “Coordination of Rab8 and Rab11 in Primary Ciliogenesis.” Proceedings of the National Academy of Sciences 107 (14): 6346–51. 10.1073/pnas.1002401107.

49. Koike-Kumagai, Makiko, Kei-ichiro Yasunaga, Rei Morikawa, Takahiro Kanamori, and Kazuo Emoto. 2009. “The Target of Rapamycin Complex 2 Controls Dendritic Tiling of Drosophila Sensory Neurons through the Tricornered Kinase Signalling Pathway.” The EMBO Journal 28 (24): 3879–92. 10.1038/emboj.2009.312.

50. Krämer, Rafael, Sandra Rode, and Sebastian Rumpf. 2019. “Rab11 Is Required for Neurite Pruning and Developmental Membrane Protein Degradation in *Drosophila* Sensory Neurons.” *Developmental Biology*, Single-cell branching morphogenesis, 451 (1): 68–78. 10.1016/j.ydbio.2019.03.003.

51. Kuo, Chay T., Lily Yeh Jan, and Yuh Nung Jan. 2005. “Dendrite-Specific Remodeling of Drosophila Sensory Neurons Requires Matrix Metalloproteases, Ubiquitin-Proteasome, and Ecdysone Signaling.” October 18, 2005. 10.1073/pnas.0507393102.

52. Kuo, Chay T., Sijun Zhu, Susan Younger, Lily Y. Jan, and Yuh Nung Jan. 2006. “Identification of E2/E3 Ubiquitinating Enzymes and Caspase Activity Regulating Drosophila Sensory Neuron Dendrite Pruning.” Neuron 51 (3): 283–90. 10.1016/j.neuron.2006.07.014.

53. Lee, Tzumin, Simone Marticke, Carl Sung, Steven Robinow, and Liqun Luo. 2000. “Cell-Autonomous Requirement of the USP/EcR-B Ecdysone Receptor for Mushroom Body Neuronal Remodeling in *Drosophila*.” Neuron 28 (3): 807–18. 10.1016/S0896-6273(00)00155-0.

54. Lin, Tzu, Hao-Hsiang Kao, Che-Hsuan Chou, Chih-Yu Chou, Yu-Ching Liao, and Hsiu-Hsiang Lee. 2020. “Rab11 Activation by Ik2 Kinase Is Required for Dendrite Pruning in Drosophila Sensory Neurons.” PLOS Genetics 16 (2): e1008626. 10.1371/journal.pgen.1008626.

55. Lin, Tzu, Po-Yuan Pan, Yu-Ting Lai, Kai-Wen Chiang, Hsin-Lun Hsieh, Yi-Ping Wu, Jian-Ming Ke, et al. 2015. “Spindle-F Is the Central Mediator of Ik2 Kinase-Dependent Dendrite Pruning in Drosophila Sensory Neurons.” PLOS Genetics 11 (11): e1005642. 10.1371/journal.pgen.1005642.

56. Magariños, Ana María, Bruce S. McEwen, Gabriele Flügge, and Eberhard Fuchs. 1996. “Chronic Psychosocial Stress Causes Apical Dendritic Atrophy of Hippocampal CA3 Pyramidal Neurons in Subordinate Tree Shrews.” Journal of Neuroscience 16 (10): 3534–40. 10.1523/JNEUROSCI.16-10-03534.1996.

57. McEwen, Bruce S., Carla Nasca, and Jason D. Gray. 2016. “Stress Effects on Neuronal Structure: Hippocampus, Amygdala, and Prefrontal Cortex.” Neuropsychopharmacology 41 (1): 3–23. 10.1038/npp.2015.171.

58. Mello, Craig, and Andrew Fire. 1995. “Chapter 19 DNA Transformation.” In Methods in Cell Biology, edited by Henry F. Epstein and Diane C. Shakes, 48:451–82. Cuenorhubditis Elegans: Modern Biologcal Analysis of an Organism. Academic Press. 10.1016/S0091-679X(08)61399-0.

59. Millward, Thomas A., Daniel Hess, and Brian A. Hemmings. 1999. “Ndr Protein Kinase Is Regulated by Phosphorylation on Two Conserved Sequence Motifs*.” Journal of Biological Chemistry 274 (48): 33847–50. 10.1074/jbc.274.48.33847.

60. Nagai, Tomoaki, and Kensaku Mizuno. 2014. “Multifaceted Roles of Furry Proteins in Invertebrates and Vertebrates.” The Journal of Biochemistry 155 (3): 137–46. 10.1093/jb/mvu001.

61. Natarajan, Rajalaxmi, Kara Barber, Amanda Buckley, Phillip Cho, Anuoluwapo Egbejimi, and Yogesh P. Wairkar. 2015. “Tricornered Kinase Regulates Synapse Development by Regulating the Levels of Wiskott-Aldrich Syndrome Protein.” PLOS ONE 10 (9): e0138188. 10.1371/journal.pone.0138188.

62. Nishida, Kei, Kenta Tsuchiya, Hiroyuki Obinata, Shizuka Onodera, Yu Honda, Yen-Cheng Lai, Nami Haruta, and Asako Sugimoto. 2021. “Expression Patterns and Levels of All Tubulin Isotypes Analyzed in GFP Knock-In C. Elegans Strains.” Cell Structure and Function 46 (1): 51–64. 10.1247/csf.21022.

63. Norkett, Rosalind, Urko del Castillo, Wen Lu, and Vladimir I Gelfand. 2020. “Ser/Thr Kinase Trc Controls Neurite Outgrowth in Drosophila by Modulating Microtubule-Microtubule Sliding.” Edited by Suzanne R Pfeffer and Jennifer G DeLuca. eLife 9 (February):e52009. 10.7554/eLife.52009.

64. Ogura, Monami, Tatsuya Kaminishi, Takayuki Shima, Miku Torigata, Nao Bekku, Keisuke Tabata, Satoshi Minami, et al. 2023. “Microautophagy Regulated by STK38 and GABARAPs Is Essential to Repair Lysosomes and Prevent Aging.” EMBO Reports 24 (12): e57300. 10.15252/embr.202357300.

65. Park, Seungmee, Nathaniel Noblett, Lauren Pitts, Antonio Colavita, Ann M. Wehman, Yishi Jin, and Andrew D. Chisholm. 2024. “Dopey-Dependent Regulation of Extracellular Vesicles Maintains Neuronal Morphology.” Current Biology: CB 34 (21): 4920–4933.e11. 10.1016/j.cub.2024.09.018.

66. Rehberg, Kati, Stefanie Kliche, Deniz A. Madencioglu, Marlen Thiere, Bettina Müller, Bernhard Manuel Meineke, Christian Freund, Eike Budinger, and Oliver Stork. 2014. “The Serine/Threonine Kinase Ndr2 Controls Integrin Trafficking and Integrin-Dependent Neurite Growth.” Journal of Neuroscience 34 (15): 5342–54. 10.1523/JNEUROSCI.2728-13.2014.

67. Ren, Peifeng, Chang-Su Lim, Robert Johnsen, Patrice S. Albert, David Pilgrim, and Donald L. Riddle. 1996. “Control of C. Elegans Larval Development by Neuronal Expression of a TGF-β Homolog.” Science 274 (5291): 1389–91. 10.1126/science.274.5291.1389.

68. Riccomagno, Martin M., and Alex L. Kolodkin. 2015. “Sculpting Neural Circuits by Axon and Dendrite Pruning.” Annual Review of Cell and Developmental Biology 31 (Volume 31, 2015): 779–805. 10.1146/annurev-cellbio-100913-013038.

69. Richardson, Claire E., Callista Yee, and Kang Shen. 2019. “A Hormone Receptor Pathway Cell-Autonomously Delays Neuron Morphological Aging by Suppressing Endocytosis.” PLoS Biology 17 (10): e3000452. 10.1371/journal.pbio.3000452.

70. Riddle, Donald L. 1977. “A Genetic Pathway for Dauer Larva Formation in Caenorhabditis Elegans : (Nematode, Development, Neuron).” https://mospace.umsystem.edu/xmlui/handle/10355/67163.

71. Riddle, Donald L., Margaret M. Swanson, and Patrice S. Albert. 1981. “Interacting Genes in Nematode Dauer Larva Formation.” Nature 290 (5808): 668–71. 10.1038/290668a0.

72. Roşianu, Flavia, Simeon R. Mihaylov, Noreen Eder, Antonie Martiniuc, Suzanne Claxton, Helen R. Flynn, Shamsinar Jalal, et al. 2023. “Loss of NDR1/2 Kinases Impairs Endomembrane Trafficking and Autophagy Leading to Neurodegeneration.” Life Science Alliance 6 (2): e202201712. 10.26508/lsa.202201712.

73. Rui, Menglong. 2024. “Recent Progress in Dendritic Pruning of Drosophila C4da Sensory Neurons.” Open Biology 14 (7): 240059. 10.1098/rsob.240059.

74. Rumpf, Sebastian, Neele Wolterhoff, and Svende Herzmann. 2019. “Functions of Microtubule Disassembly during Neurite Pruning.” Trends in Cell Biology 29 (4): 291–97. 10.1016/j.tcb.2019.01.002.

75. Santos, Paulo F., Beatriz Fazendeiro, Francis C. Luca, António Francisco Ambrósio, and Hélène Léger. 2023. “The NDR/LATS Protein Kinases in Neurobiology: Key Regulators of Cell Proliferation, Differentiation and Migration in the Ocular and Central Nervous System.” European Journal of Cell Biology 102 (2): 151333. 10.1016/j.ejcb.2023.151333.

76. Schroeder, Nathan E., Rebecca J. Androwski, Alina Rashid, Harksun Lee, Junho Lee, and Maureen M. Barr. 2013. “Dauer-Specific Dendrite Arborization in C. Elegans Is Regulated by KPC-1/Furin.” Current Biology: CB 23 (16): 1527–35. 10.1016/j.cub.2013.06.058.

77. Schuldiner, Oren, and Avraham Yaron. 2015. “Mechanisms of Developmental Neurite Pruning.” Cellular and Molecular Life Sciences 72 (1): 101–19. 10.1007/s00018-014-1729-6.

78. Shi, Rebecca, Xue Yan Ho, Li Tao, Caitlin A. Taylor, Ting Zhao, Wei Zou, Malcolm Lizzappi, Kelsie Eichel, and Kang Shen. 2024. “Stochastic Growth and Selective Stabilization Generate Stereotyped Dendritic Arbors.” bioRxiv. 10.1101/2024.05.08.591205.

79. Smith, Cody J., Joseph D. Watson, W. Clay Spencer, Tim O’Brien, Byeong Cha, Adi Albeg, Millet Treinin, and David M. Miller. 2010. “Time-Lapse Imaging and Cell-Specific Expression Profiling Reveal Dynamic Branching and Molecular Determinants of a Multi-Dendritic Nociceptor in *C. Elegans*.” Developmental Biology 345 (1): 18–33. 10.1016/j.ydbio.2010.05.502.

80. Suzuki, Atsushi, Tsutomu Ogura, and Hiroyasu Esumi. 2006. “NDR2 Acts as the Upstream Kinase of ARK5 during Insulin-like Growth Factor-1 Signaling.” Journal of Biological Chemistry 281 (20): 13915–21. 10.1074/jbc.M511354200.

81. Tamaskovic, Rastislav, Samuel J. Bichsel, and Brian A. Hemmings. 2003. “NDR Family of AGC Kinases – Essential Regulators of the Cell Cycle and Morphogenesis.” FEBS Letters 546 (1): 73–80. 10.1016/S0014-5793(03)00474-5.

82. Tamaskovic, Rastislav, Samuel J. Bichsel, Hélène Rogniaux, Mario R. Stegert, and Brian A. Hemmings. 2003. “Mechanism of Ca2+-Mediated Regulation of NDR Protein Kinase through Autophosphorylation and Phosphorylation by an Upstream Kinase*.” Journal of Biological Chemistry 278 (9): 6710–18. 10.1074/jbc.M210590200.

83. Taylor, Caitlin A., Jing Yan, Audrey S. Howell, Xintong Dong, and Kang Shen. 2015. “RAB-10 Regulates Dendritic Branching by Balancing Dendritic Transport.” PLOS Genetics 11 (12): e1005695. 10.1371/journal.pgen.1005695.

84. Ultanir, Sila K., Nicholas T. Hertz, Guangnan Li, Woo-Ping Ge, Alma L. Burlingame, Samuel J. Pleasure, Kevan M. Shokat, Lily Yeh Jan, and Yuh-Nung Jan. 2012. “Chemical Genetic Identification of NDR1/2 Kinase Substrates AAK1 and Rabin8 Uncovers Their Roles in Dendrite Arborization and Spine Development.” Neuron 73 (6): 1127–42. 10.1016/j.neuron.2012.01.019.

85. Verde, Fulvia, David J. Wiley, and Paul Nurse. 1998. “Fission Yeast Orb6, a Ser/Thr Protein Kinase Related to Mammalian Rho Kinase and Myotonic Dystrophy Kinase, Is Required for Maintenance of Cell Polarity and Coordinates Cell Morphogenesis with the Cell Cycle.” June 1998. 10.1073/pnas.95.13.7526.

86. Vyas, Ajai, Rupshi Mitra, B. S. Shankaranarayana Rao, and Sumantra Chattarji. 2002. “Chronic Stress Induces Contrasting Patterns of Dendritic Remodeling in Hippocampal and Amygdaloid Neurons.” Journal of Neuroscience 22 (15): 6810–18. 10.1523/JNEUROSCI.22-15-06810.2002.

87. Westlake, Christopher J., Lisa M. Baye, Maxence V. Nachury, Kevin J. Wright, Karen E. Ervin, Lilian Phu, Cecile Chalouni, et al. 2011. “Primary Cilia Membrane Assembly Is Initiated by Rab11 and Transport Protein Particle II (TRAPPII) Complex-Dependent Trafficking of Rabin8 to the Centrosome.” Proceedings of the National Academy of Sciences of the United States of America 108 (7): 2759–64. 10.1073/pnas.1018823108.

88. White, J. G., E. Southgate, J. N. Thomson, and S. Brenner. 1986. “The Structure of the Nervous System of the Nematode Caenorhabditis Elegans.” *Philosophical Transactions of the Royal Society of London. Series B*, Biological Sciences 314 (1165): 1–340.

89. Williams, Darren W., and James W. Truman. 2005. “Cellular Mechanisms of Dendrite Pruning in Drosophila: Insights from in Vivo Time-Lapse of Remodeling Dendritic Arborizing Sensory Neurons.” Development 132 (16): 3631–42. 10.1242/dev.01928.

90. Wu, Zhihao, Tomoyo Sawada, Kahori Shiba, Song Liu, Tomoko Kanao, Ryosuke Takahashi, Nobutaka Hattori, Yuzuru Imai, and Bingwei Lu. 2013. “Tricornered/NDR Kinase Signaling Mediates PINK1-Directed Mitochondrial Quality Control and Tissue Maintenance.” Genes & Development 27 (2): 157–62. 10.1101/gad.203406.112.

91. Yd, Tay, Leda M, Spanos C, Rappsilber J, Goryachev Ab, and Sawin Ke. 2019. “Fission Yeast NDR/LATS Kinase Orb6 Regulates Exocytosis via Phosphorylation of the Exocyst Complex.” Cell Reports 26 (6). 10.1016/j.celrep.2019.01.027.

92. Zallen, Jennifer A., Erin L. Peckol, David M. Tobin, and Cornelia I. Bargmann. 2000. “Neuronal Cell Shape and Neurite Initiation Are Regulated by the Ndr Kinase SAX-1, a Member of the Orb6/COT-1/Warts Serine/Threonine Kinase Family.” Molecular Biology of the Cell 11 (9): 3177–90. 10.1091/mbc.11.9.3177.

93. Zhang, Heng, Yan Wang, Jack Jing Lin Wong, Kah-Leong Lim, Yih-Cherng Liou, Hongyan Wang, and Fengwei Yu. 2014. “Endocytic Pathways Downregulate the L1-Type Cell Adhesion Molecule Neuroglian to Promote Dendrite Pruning in Drosophila.” Developmental Cell 30 (4): 463–78. 10.1016/j.devcel.2014.06.014.

94. Zhang, Lei, Fengyuan Tang, Luigi Terracciano, Debby Hynx, Reto Kohler, Sandrine Bichet, Daniel Hess, et al. 2015. “NDR Functions as a Physiological YAP1 Kinase in the Intestinal Epithelium.” Current Biology: CB 25 (3): 296–305. 10.1016/j.cub.2014.11.054.

95. Zou, Wei, Smita Yadav, Laura DeVault, Yuh Nung Jan, and David R. Sherwood. 2015. “RAB-10-Dependent Membrane Transport Is Required for Dendrite Arborization.” PLOS Genetics 11 (9): e1005484. 10.1371/journal.pgen.1005484.

